# Human non-canonical inflammasomes activate CASP3 to limit intracellular *Salmonella* replication in macrophages

**DOI:** 10.1101/2025.10.31.685927

**Authors:** Madhura Kulkarni, Christopher M. Bourne, Ashutosh B. Mahale, Patrick M. Exconde, Cecelia Murphy, Sofia Cervantes, Matilda Kardhashi, Mirai Kambayashi, William Yoo, Tristan J. Wrong, Robert C. Patio, Bohdana M. Discher, Cornelius Y. Taabazuing

## Abstract

Inflammasomes are multiprotein signaling platforms that activate inflammatory caspases to initiate innate immune signaling. In humans, canonical inflammasomes activate CASP1, which cleaves the pore-forming protein gasdermin D (GSDMD) and the cytokines IL-1β and IL-18. In contrast, the non-canonical inflammasome detects bacterial lipopolysaccharide (LPS) through CASP4/5, which cleave GSDMD to drive pyroptosis. While CASP1 substrates are well characterized, CASP4/5 substrates remain less defined. Here, we show that in response to intracellular LPS and gram-negative bacterial infection, CASP4/5 directly cleave and activate the executioner caspases CASP3/7. CASP3 in turn cleaves and activates gasdermin E (GSDME). Surprisingly, CASP3, but not GSDME, was required for restricting intracellular *Salmonella* replication, suggesting that CASP4/5-induced apoptosis contributes to host defense. We further show that most GSDMD cleavage during non-canonical inflammasome activation is mediated by CASP1, and that GSDMD is the primary driver of pyroptosis. Finally, we confirm that CASP4/5 activate CASP3/7 and GSDME in human primary macrophages. These findings establish CASP4/5 as dual apoptotic initiator and inflammatory caspases and reveal a central role for the apoptotic signaling cascade in non-canonical inflammasome-mediated immunity.

## Introduction

The detection and elimination of invading pathogens by the immune system is essential for host survival. It is critical that the immune response to invading pathogens be tightly regulated such that an appropriate response can be mounted without excessive inflammation, a condition known as sepsis. Sepsis is a life threatening condition which mainly occurs in children under the age of five and is one of the leading causes of death worldwide^1,2^.

A key component of the host response to invading pathogens is the assembly of multi-protein signaling complexes called inflammasomes^3^. Inflammasome assembly activates cysteine proteases called caspases that induce inflammation and lytic cell death known as pyroptosis, which is critical for infection control^4,5^. The canonical inflammasome pathway activates caspase-1 (CASP1), while the non-canonical pathway activates caspase-4 and caspase-5 (CASP4/5) in humans and caspase-11 (CASP11) in mice, collectively referred to as inflammatory caspases^6,7^. CASP1 activation can be triggered by diverse stimuli, such as host or pathogen proteases, and potassium efflux^8-11^. In contrast, CASP4/5 and -11 sense and are activated by intracellular lipopolysaccharide (LPS) from Gram-negative bacteria like *Salmonella enterica* serovar Typhimurium (*Stm*), an intracellular pathogen that is a major cause of food-borne disease globally^6,12-14^. A recent study demonstrated that while CASP1 is important during early infection, CASP4 is essential in preventing intracellular *Salmonella* replication at late stages of infection in human macrophages^15^. However, the mechanism by which CASP4 restricts replication at late stages remains unclear.

The substrates of inflammatory caspases are crucial mediators of immunity, yet their full repertoire and functional significance remain incompletely defined^16^. Human inflammatory caspases are known to cleave the interleukin family of cytokines, IL-1β and IL-18, and the pore-forming protein gasdermin D (GSDMD), leading to pyroptosis^17-23^. In this manner, pyroptosis protects the host from infection^24-28^. In the absence of GSDMD, CASP1 can process the apoptotic executioner caspases-3 and -7 (CASP3/7) to induce apoptosis^29^. A proteomic screen also identified CASP7 as a substrate of inflammasome-activated CASP1 in mice, but in the same study, CASP3 activation was found to be independent of CASP1^30^. It was later shown that, like human cells, CASP1 could mediate CASP3 processing in mice to induce apoptosis in the absence of GSDMD when infected with *Salmonella enterica* serovar Typhimurium or poly(dA:dT)^31,32^. This process was dependent on CASP1 cleaving Bid (BH3-interacting domain death agonist), which subsequently induced caspase-9 (CASP9)-dependent activation of CASP3^31,32^. These findings suggest that CASP1 activation can lead to either direct or indirect activation of executioner CASP3 and -7 and that inflammatory caspases can initiate apoptosis.

Apoptosis is a regulated form of cell death that maintains organismal development and homeostasis and is generally thought to be immunologically silent. Emerging evidence suggests that the induction of apoptosis during infections may have important consequences for host defense. For example, dendritic cells activate both pyroptosis and apoptosis during late stages of infection and this is essential for restricting *Legionella pneumophila* replication^33,34^. Similarly, activation of the apoptotic signaling cascade during viral infection led to CASP3-dependent cleavage of gasdermin E (GSDME), which induced pyroptosis and protected human skin organoids from viral replication^35,36^. Whether the induction of apoptosis or CASP3 and GSDME-mediated pyroptosis contributes broadly to host defense remains unknown. Moreover, while CASP1 is well-established to activate the apoptotic signaling cascade, whether human non-canonical inflammasomes (CASP4/5) similarly cleave apoptotic caspases remains less clear.

Substrates of the non-canonical inflammasome pathway are generally not well-defined. A recent proteomics screen identified CASP7 as a substrate of CASP4 in humans, though its functional significance remained unknown^37^. We also recently reported that IL-1β and IL-18 are substrates of CASP4/5^21^. Notably, in CASP4/5-deficient cells, we observed a loss of CASP3 activation in response to intracellular LPS, suggesting that the human non-canonical inflammasomes engage the apoptotic pathway. However, whether this is a direct processing event and its functional significance regarding pathogen defense remained to be elucidated.

In this study, we identify CASP3 and CASP7 as direct substrates of human inflammatory caspases. We found that CASP4/5 directly cleave CASP3/7, and active CASP3 subsequently cleaves and activates GSDME during intracellular LPS stimulation and *Salmonella* infection. CASP3-induced GSDME-mediated pyroptosis has been described in the context of chemotherapeutics activating apoptotic signaling or during cytotoxic lymphocyte-mediated cell death^38^, but this pathway has not been known to be activated during non-canonical inflammasome activation. We demonstrate that CASP3/7 and GSDME processing occurs in primary human cells in response to intracellular LPS and *Salmonella* infection. Interestingly, we discovered that GSDME was dispensable, but CASP3 was required for inhibiting intracellular *Salmonella* replication, demonstrating that non-canonical inflammasomes mediate host defense in part by activating the apoptotic signaling cascade.

## Results

### Human inflammatory caspases directly cleave and activate executioner CASP3 and CASP7

Because previous work has shown that CASP1 can cleave CASP3/7, and CASP4 has been implicated in CASP7 processing^37^, we first investigated whether inflammatory caspases directly target CASP3 and CASP7. We expressed human GSDMD (as a control), CASP3, or CASP7 in HEK 293T cells and incubated lysates with recombinant active CASP1, CASP4, CASP5, CASP8, or mouse CASP11 for 1 or 3 hours. As expected, inflammatory caspases efficiently processed GSDMD at both time points (Fig. 1A). CASP1, CASP4, CASP5 and CASP8 processed human CASP3 (Fig. 1B). Mouse CASP11 treated lysates exhibited very minimal processing of human CASP3 at prolonged incubation, suggesting a divergence between human and murine non-canonical inflammasomes. This aligns with prior reports that CASP4/5 can cleave human IL-18, whereas CASP11 cannot cleave mouse IL-18^20-22^. Surprisingly, only CASP1 and CASP4 processed CASP7 in this assay (Fig. 1C), despite the well-established role of CASP8 in cleaving both CASP7 and CASP3^39^ (Fig. 1B,C), indicating that complementary methods are needed to confirm processing.

**Figure 1.**
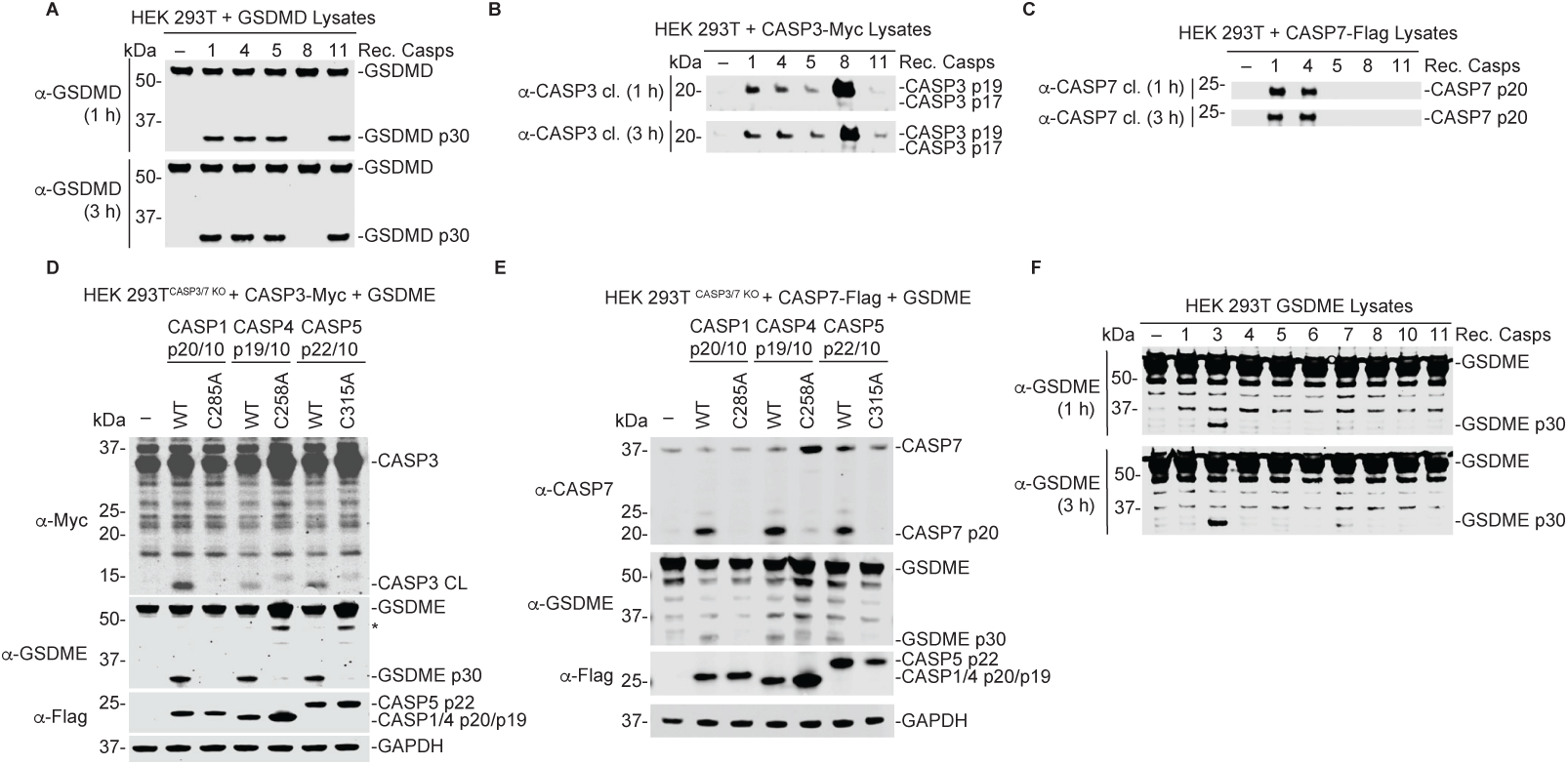
Caspases-1, -4, and -5 directly cleave and activate caspases-3 and -7. (A-C) HEK 293T cells were transiently transfected with plasmids coding for human GSDMD **(A)**, CASP3 **(B)**, or CASP7 **(C)**. Lysates were then incubated with 0.25 activity units/μL of each indicated recombinant caspase for 1 or 3 h and cleavage products were assessed by immunoblotting. **(D,E)** *CASP3/7* KO HEK 293T cells were transiently co-transfected with the indicated constructs coding for active CASP3 **(D)** or CASP7 **(E)** along with GSDME, and catalytically active or inactive CASP1, -4, or -5 for 24 h before the cell lysates were analyzed by immunoblotting. **(F)** HEK 293T cells were transiently transfected with a plasmid coding for GSDME for 24 h. Lysates were harvested and incubated with 0.25 activity units/μL of each recombinant caspase for 1 or 3 h. Cleavage products were assessed by immunoblotting. Data are representative of three or more independent experiments.

We next sought to determine whether active inflammatory caspases would cleave CASP3/7 in cells. We generated *CASP3/7* double knockout (KO) HEK 293T cells (Fig. S1A) and co-transfected these cells with either active or catalytically inactive human inflammatory caspases along with catalytically inactive CASP3 or CASP7. Consistent with our *in vitro* assays, all inflammatory caspases processed catalytically inactive CASP3 and CASP7 (Fig. S1B,C), and this processing was dependent on their catalytic activity and not on the catalytic activity of CASP3 or CASP7.

We also confirmed that inflammatory caspases cleave CASP3/7 using an orthogonal activation mechanism. We utilized our previously established dimerizable inflammatory caspase cell system in which cells express a modified caspase where the caspase activation and recruitment domains (CARD) are swapped with a dimerizable domain (DmrB), allowing selective activation of CASP1, CASP4, or CASP5 by the small molecule drug AP20187^21^. HEK 293T cells stably expressing both a DmrB-caspase and IL-18, a known substrate of inflammatory caspases that serves as a positive control, were transfected with either CASP3 or CASP7. Following AP20187 treatment, we observed activation of the DmrB-CASP1, -4, and -5, as indicated by IL-18 cleavage (Fig. S1D-I). Consistent with our *in vitro* biochemical assays, activated CASP1, -4, and -5 all processed CASP3 (Fig. S1D-F) and CASP7 (Fig. S1G-I). These findings suggest that upon direct activation in cells, inflammatory caspases cleave the executioner caspases CASP3 and CASP7.

Caspases cleave peptide bonds at aspartic (Asp) acid residues^40,41^. During apoptosis, the initiator caspases (CASP8/9) activate executioner caspases by cleaving CASP3 at Asp175 and CASP7 at Asp198, generating catalytically active large and small subunits^30,42^. Because inflammatory caspase processing of CASP3/7 produced fragments that were detected by antibodies specific for their active forms, we hypothesized that cleavage activates the executioner caspases. To test this, we co-expressed catalytically competent CASP3 (Fig. 1D) or CASP7 (Fig. 1E) with GSDME, a known substrate of CASP3^43^, along with either catalytically active or inactive human inflammatory caspases in HEK 293T *CASP3/7* KO cells. Consistent with their activation, CASP3, and to a lesser extent CASP7, induced GSDME cleavage, indicating that inflammatory caspases cleave and activate CASP3/7. We ruled out that inflammatory caspases do not directly process GSDME through recombinant protein assays. Lysates from GSDME-expressing HEK293T cells were incubated with apoptotic (CASP3, CASP6, CASP7, CASP8) or inflammatory (CASP1, CASP4, CASP5, CASP11) caspases (Fig. 1C). As expected, CASP3 robustly cleaved GSDME, while CASP7 produced only minimal processing after prolonged incubation, suggesting CASP7-mediated GSDME cleavage is possible but likely not physiologically relevant. Collectively, these findings support that human CASP1/4/5 directly cleave CASP3/7 and promote their activation in cellular and lysate systems, with CASP3 as the primary mediator of GSDME cleavage.

### CASP1, CASP8, and CASP9 are dispensable for CASP3/7 and GSDME processing during non-canonical inflammasome activation by intracellular LPS

Our in vitro and cellular assays indicated that inflammatory caspases cleave and activate CASP3/7, and that CASP3 can subsequently cleave GSDME to drive pyroptosis^38^. Although CASP3-mediated GSDME cleavage has been shown to induce pyroptosis, whether GSDME contributes to non-canonical inflammasome mediated cell death is unknown. To address this, we transfected LPS into PMA-differentiated THP-1 macrophages to activate CASP4/5 (Fig. 2A). CASP1 can be activated via the NLRP3 inflammasome downstream of CASP4/5 due to K+ efflux through GSDMD pores^8,9,44^. Prior work indicated that during CASP1 activation, both GSDME and GSDMD contribute to cell death and cytokine release^45^. To rule out CASP1-mediated cleavage of CASP3/7 downstream of CASP4/5, we pretreated THP1 cells with the NLRP3 inhibitor MCC950^46^ prior to LPS transfection (Fig. 2A). LPS treatment resulted in CASP3, CASP7, and GSDME cleavage. As expected, LPS also induced robust GSDMD cleavage and CASP1 activation, as evidenced by the appearance of the cleaved 30 kDa fragment of GSDMD and p10 species of CASP1 by immunoblotting (Fig. 2A). MCC950 abolished CASP1 activation, which led to a slight decrease in GSDMD processing (Fig. 2A). Interestingly, we observed increased CASP3, CASP7 and GSDME cleavage in MCC950-treated cells compared to LPS-only treated cells (Fig. 2A), suggesting that in the absence of CASP1, CASP4/5 activation induces robust processing and activation of CASP3/7, and GSDME. Consistent with this notion and the increased CASP3/7 processing, we observed increased CASP3/7 activity compared to control and LPS-only treated cells (Fig. 2C) using an intracellular CASP3/7 activity reporter dye. Despite loss of CASP1 activation, the kinetics of Sytox Green uptake remained unchanged in MCC950-treated cells compared to LPS-only treated cells, likely due to elevated GSDME processing (Fig. 2B).

**Figure 2.**
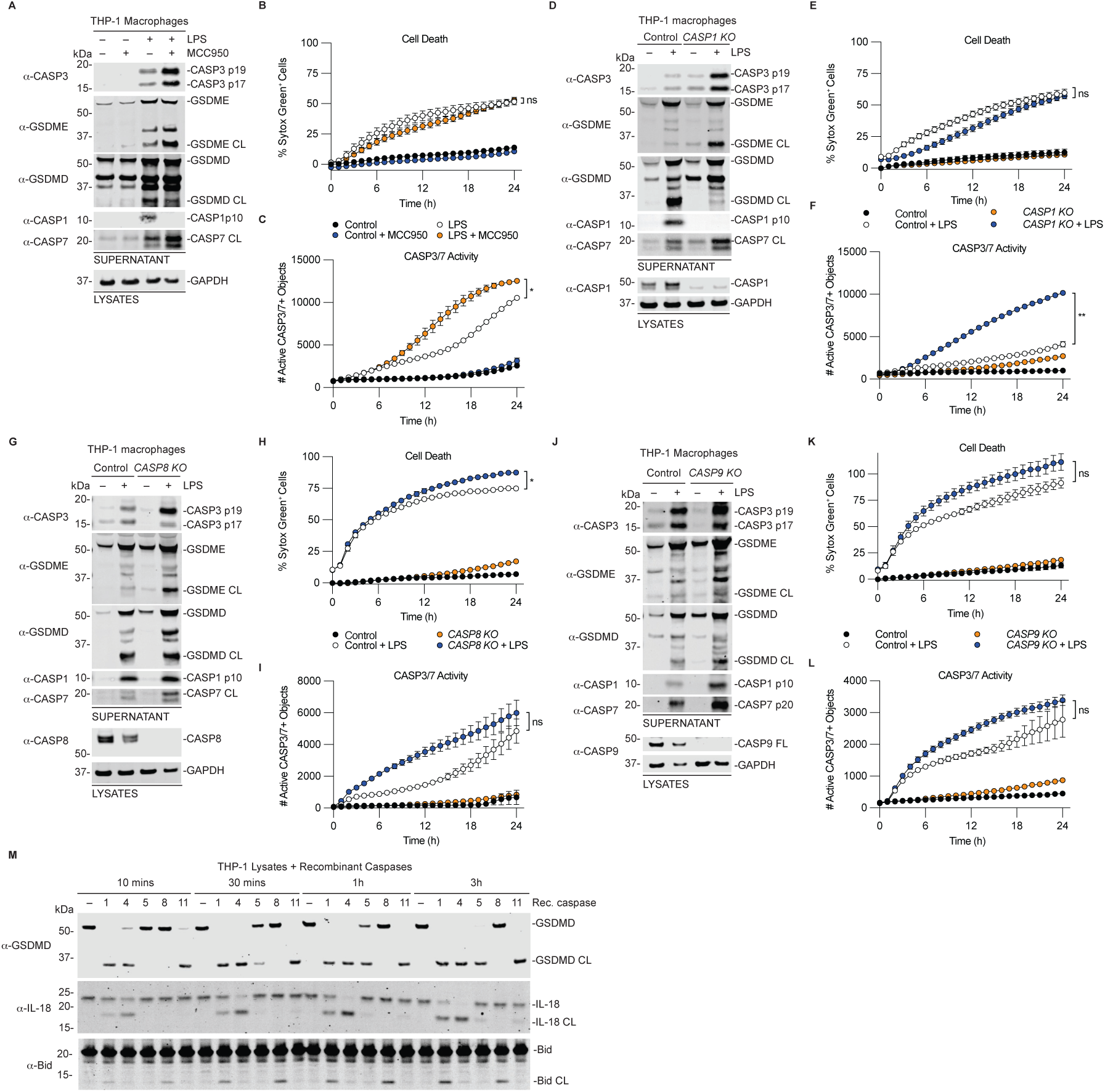
CASP1, CASP8, and CASP9 are dispensable for CASP3/7 and GSDME processing during non-canonical inflammasome activation by intracellular LPS. (**A-C**) WT THP-1 cells were transfected with 25 μg/mL LPS in the presence or absence of the NLRP3 inhibitor MCC950 (10 µM) for 24 h. Supernatants and lysates were then analyzed by immunoblotting (**A**), cell death was measured by monitoring Sytox Green uptake (**B**), and CASP3/7 activity was determined using CellTreat CASP3/7 detection reagent (**C**). (**D-F**) Control-matched *CASP1* KO (**D-F**), *CASP8* KO (**G-I**), and *CASP9* KO (**J-L**) THP-1 cells were transfected with 25 μg/mL LPS for 24 h then supernatants and lysates were then analyzed by immunoblotting (**D, G, J**), cell death was measured by monitoring Sytox Green uptake (**E, H, K**), and CASP3/7 activity was determined using CellTreat CASP3/7 detection reagent (**F, I, L**). (**M**) PMA-differentiated WT THP-1 macrophages were lysed and incubated with 0.25 activity units/μL of recombinant caspases for the indicated times before immunoblot analysis. Data are means ± SEM of three independent replicates, and representative of at least three independent experiments except for M (n = 2). ****P < 0.0001, ***P < 0.001, **P < 0.01, and *P < 0.05 by two-way ANOVA test with Tukey’s multiple comparison test comparing the control treated to KO treated samples at 24 h.

Because GSDME can be processed by CASP3 downstream of the extrinsic and intrinsic apoptotic pathways mediated by CASP8 and CASP9 respectively^43^, we next sought to genetically dissect the contribution of CASP1, CASP8, and CASP9 to CASP3 activation and GSDME processing. In *CASP1* KO THP-1 cells, LPS still triggered robust cleavage of CASP3, CASP7, and GSDME, accompanied by increased CASP3/7 activity (Fig. 2D-F). In contrast, GSDMD processing was nearly abolished, implying that most GSDMD cleavage downstream of non-canonical inflammasome activation is driven by CASP1. This aligns with our previous observations that in cells, GSDMD is a poor substrate for CASP4 and -5 compared to CASP1^21^. Importantly, these results indicate that CASP1 is not required for CASP3/7, and GSDME processing.

Both CASP8 and CASP9 are known to activate CASP3 and CASP7 to induce apoptosis, a process constantly ongoing to maintain cellular homeostasis^47,48^. Like in *CASP1* KO cells, deletion of CASP8 (Fig. 2G-I) or CASP9 (Fig. 2J-L) did not block CASP3/7 or GSDME processing, nor alter GSDMD cleavage or pyroptosis, as assessed by Sytox Green uptake. In fact, CASP3/7 activity was modestly increased, suggesting that CASP8 or CASP9 may compete with CASP4/5 for CASP3/7 engagement.

Finally, because CASP1 is known to cleave Bid, a member of the Bcl-2 family of proteins that activates CASP9 to induce CASP3/7 cleavage^31,32,49^, we sought to determine whether CASP4/5 activate CASP3/7 indirectly through activation of Bid. CASP3 is also known to cleave Bid^50^. To rule out a contribution from CASP3 activated by CASP4/5 in cells, we incubated lysates from THP-1 macrophages with equivalent activity units of recombinant caspases and assessed processing of substrates at early time points preceding CASP3/7 activation by immunoblotting.

As expected, CASP1 and CASP8 processed Bid into the truncated form (Fig. 2M). However, CASP4 and CASP5 failed to cleave Bid, but were active and able to process IL-18 and GSDMD (Fig. 2M). In this assay, where multiple substrates are present simultaneously, IL-18 processing appeared to be favored by CASP4 compared to other caspases while GSDMD processing was favored by CASP1 (Fig. 2M). Taken together, these data indicate that CASP1, CASP8, and CASP9 are dispensable for CASP3/7 activation during noncanonical inflammasome activation and support a model in which CASP4/5 directly cleave CASP3/7 to promote CASP3/GSDME-mediated pyroptosis.

### CASP3, and CASP4/5 are required to induce GSDME processing during non-canonical inflammasome activation by intracellular LPS

Our findings above suggest that CASP4/5 activate the executioner CASP3/7, leading to CASP3/GSDME-mediated pyroptosis. To directly test this, we transfected WT, *CASP4/5* KO, *CASP3* KO, or *CASP7* KO cells with LPS (Fig. 3A-I). We then assessed CASP3/7 and GSDME processing by immunoblotting, cell death by Sytox Green uptake, and CASP3/7 activity using a cell permeable reporter dye. Loss of CASP4/5 significantly impaired CASP3/7 and GSDME cleavage, cell death, and CASP3/7 activity compared to control cells in response to intracellular LPS (Fig 3A-C). *CASP3* KO cells lacked GSDME cleavage but exhibited increased GSDMD processing in LPS transfected cells (Fig. 3D). Despite loss of GSDME processing, pyroptosis was not significantly diminished (Fig. 3E), suggesting functional compensation through enhanced GSDMD activation. CASP3 deficiency slightly reduced overall CASP3/7 activity, likely due to preserved CASP7 activity (Fig. 3F). Unexpectedly, *CASP7* KO cells retained CASP3 activation and GSDME processing (albeit less than control cells), but CASP1 processing into the p10 species and GSDMD processing were nearly abolished, resulting in delayed cell death (Fig. 3G,H). *CASP7* KO cells also had attenuated CASP3/7 activity, as expected (Fig. 3I). Taken together, these findings demonstrate that CASP4/5 activate CASP3/7, with CASP3 serving as the primary protease that cleaves GSDME. While CASP3/GSDME activation can drive pyroptosis downstream of the non-canonical inflammasome in part, loss of CASP3 is buffered by increased GSDMD activity, highlighting plasticity in the execution of pyroptosis.

**Figure 3.**
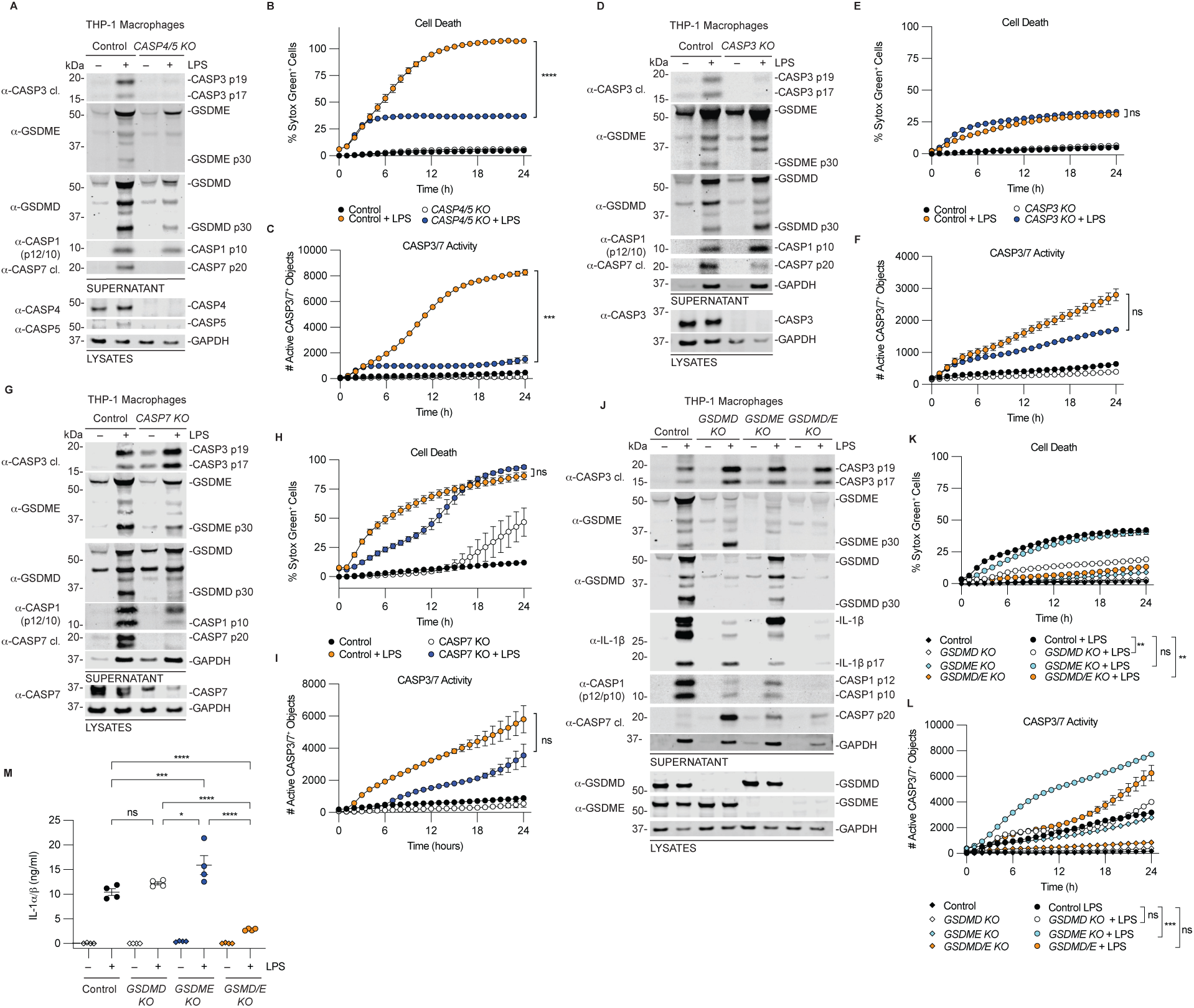
CASP3, and CASP4/5 are required to induce GSDME processing during non-canonical inflammasome activation by intracellular LPS. (**A-C**) WT and *CASP4/5* double KO THP-1 cells were transfected with 25 μg/mL of LPS for 24 h then supernatants and lysates were analyzed by immunoblotting (**A**), cell death was measured by monitoring Sytox Green uptake (**B**), and CASP3/7 activity was determined using CellTreat CASP3/7 detection reagent (**C**). (**D-F**) WT and *CASP3* KO THP-1 cells were transfected with 25 μg/mL of LPS for 24 h then supernatants and lysates were then analyzed by immunoblotting (**H**), cell death was measured by monitoring Sytox Green uptake (**I**), and CASP3/7 activity was determined using CellTreat CASP3/7 detection reagent. (**G-I**) WT and *CASP7* KO THP-1 cells were transfected with 25 μg/mL of LPS for 24 h then supernatants and lysates were then analyzed by immunoblotting (**H**), cell death was measured by monitoring Sytox Green uptake (**I**), and CASP3/7 activity was determined using CellTreat CASP3/7 detection reagent. (**J-M**) WT, *GSDMD* KO, *GSDME* KO and *GSDMD/E* KO THP-1 macrophages were transfected with 25 μg/mL of LPS for 24 h then supernatants and lysates were analyzed by immunoblotting (**J**), cell death was measured by Sytox Green uptake (**K**), and CASP3/7 activity was determined using CellTreat CASP3/7 detection reagent (**L**), and the amount of active IL-α/ý released into the supernatants was quantified (**M**). Data are means ± SEM of three independent replicates, and representative of at least three independent experiments. ****P < 0.0001, ***P < 0.001, **P < 0.01, and *P < 0.05 by two-way ANOVA test with Tukey’s multiple comparison test comparing the control treated to KO treated samples at 24 h. Data in M were analyzed by One-way Anova with Tukey’s multiple comparisons test.

Mammals encode six gasdermins: gasdermin A, B, C, D, E, and PJVK (also known as DFNB59)^36,51,52^. Typically, their N-terminal pore-forming domains are autoinhibited by the C-termini, but upon cleavage, the N-termini are liberated and form pores to induce pyroptosis. Inflammatory caspases cleave GSDMD to induce pyroptosis^17-19^. While CASP3 has been reported to cleave GSDME to induce pyroptosis, this has primarily been described in the context of chemotherapy-induced apoptosis or cytotoxic lymphocyte-mediated cell death^38,53,54^. Whether CASP3/GSDME contributes to pyroptosis during non-canonical inflammasome activation has not been established.

To address this, we transfected WT, *GSDMD* KO, *GSDME* KO, or *GSDMD/E* double KO THP-1 macrophages with LPS and assessed gasdermin processing, cell death, CASP3/7 activation, and IL-1α/β release (Fig. 3J-M). Immunoblots revealed increased CASP3/7 processing, but decreased CASP1 processing in *GSDMD*, *GSDME*, or *GSDMD/E* KO cells compared to control cells with LPS transfection (Fig. 3J), suggesting that both GSDMD and GSDME may contribute to downstream CASP1 activation via the NLRP3 inflammasome. Loss of either GSDMD or GSDME led to increased CASP3 and CASP7 activation compared to WT cells. However, the CASP3/7 activity was only significantly increased in *GSDME* KO cells (Fig. 3L). GSDME processing was increased in *GSDMD* KO cells, but GSDMD processing was attenuated in *GSDME* KO cells compared to WT cells (Fig. 3J). Interestingly, GSDME loss did not affect Sytox Green uptake, whereas GSDMD deficiency markedly reduced Sytox Green uptake, and *GSDMD/E* double KO cells phenocopied *GSDMD* KO cells (Fig. 3K), implying that GSDMD is the principal driver of lytic cell death in response to intracellular LPS.

In contrast, IL-1α/ý release required both gasdermins, as assessed by the HEK Blue assay which reports on the activity of active IL-1α/ý^21^ (Fig. 3M). Either GSDMD or GSDME was sufficient to release similar amounts of active IL-1α/ý as WT cells, but loss of both GSDMD/E resulted in significantly attenuated cytokine release (Fig. 3M), consistent with reports that both contribute to cytokine release during canonical inflammasome signaling^45^. Altogether these findings suggest that both GSDMD and GSDME contribute to cytokine release during non-canonical inflammasome activation, but GSDMD is the primary driver of pyroptosis.

### CASP3, and CASP4/5 are required for GSDME processing during non-canonical inflammasome activation by intracellular *Salmonella*

The non-canonical inflammasomes detect LPS from intracellular gram-negative bacteria. We next asked if CASP4/5 cleave and activate CASP3/7 during bacterial infections. We infected WT, *CASP4/5* KO, and *CASP3* KO cells with *Salmonella enterica* serovar Typhimurium (*Stm*) then assessed CASP3/7 and GSDME processing by immunoblotting, cell death by Sytox Green uptake, and CASP3/7 activity using a cell permeable reporter dye. Log-phase *Salmonella* induces expression of the type 3 secretion system (T3SS) encoded by *Salmonella* pathogenicity island 1 (SPI-1) that activates both canonical and non-canonical inflammasome signaling ^55-58^. Thus, we first infected WT and *CASP4/5* KO (Fig. S2A,B) or *CASP3* KO (Fig. S2C, D) THP1-1 macrophages with *Salmonella* grown to stationary phase (*Stm*^Stat.^) to minimize the contribution of the canonical caspase-1 signaling pathway then measured cell death and processing of caspases and gasdermins by immunoblotting. Since CASP4 is thought to contribute to resolving infection at later time in infection^15^, we tested different multiplicity of infections (MOIs) to mimic different infection burdens. In control cells, we observed dose-dependent cleavage of GSDME and GSDMD (Fig. S2A,C). Importantly, CASP3/7 and GSDME processing was largely abrogated in *CASP4/5* KO cells while GSDMD processing was retained, likely due to the activation of the NLRP3 inflammasomes (Fig. S2A)^58-60^. GSDME, but not GSDMD processing, was also abrogated in *CASP3* KO cells (Fig. S2C). Interestingly, despite cleavage of the pore-forming proteins, we observed little to no cell death by Sytox Green (Fig. S2B,D). This suggests that CASP3/7 activation and subsequent GSDME cleavage is CASP4/5-dependent, but processing does not generate enough pores to induce cell lysis with stationary phase *Salmonella*.

Given that there was little cell death observed with (*Stm*^Stat.^) and the fact that SPI-1 is important for *Salmonella* to invade and replicate in host cells^61,62^, we focused on SPI-1-induced infection models for further studies. We infected WT, *CASP4/5* KO and *CASP3* KO cells with SPI-1-induced *Salmonella* (*Stm*^SPI-1^) (Fig. 4A-D). Although CASP1 was activated and could contribute to CASP3 activation, CASP3/7 activation and CASP3-dependent processing of GSDME were predominantly mediated by CASP4/5 (Fig. 4A,C). However, consistent with the notion that CASP1 is the dominant protease that processes GSDMD in cells, SPI-1-induced *Stm* infection triggered more cell death than stationary-phase *Stm* infections, which was attenuated in *CASP4/5* KO cells compared to control cells, but unchanged between control and *CASP3* KO cells (Fig. 4B,D).

**Figure 4.**
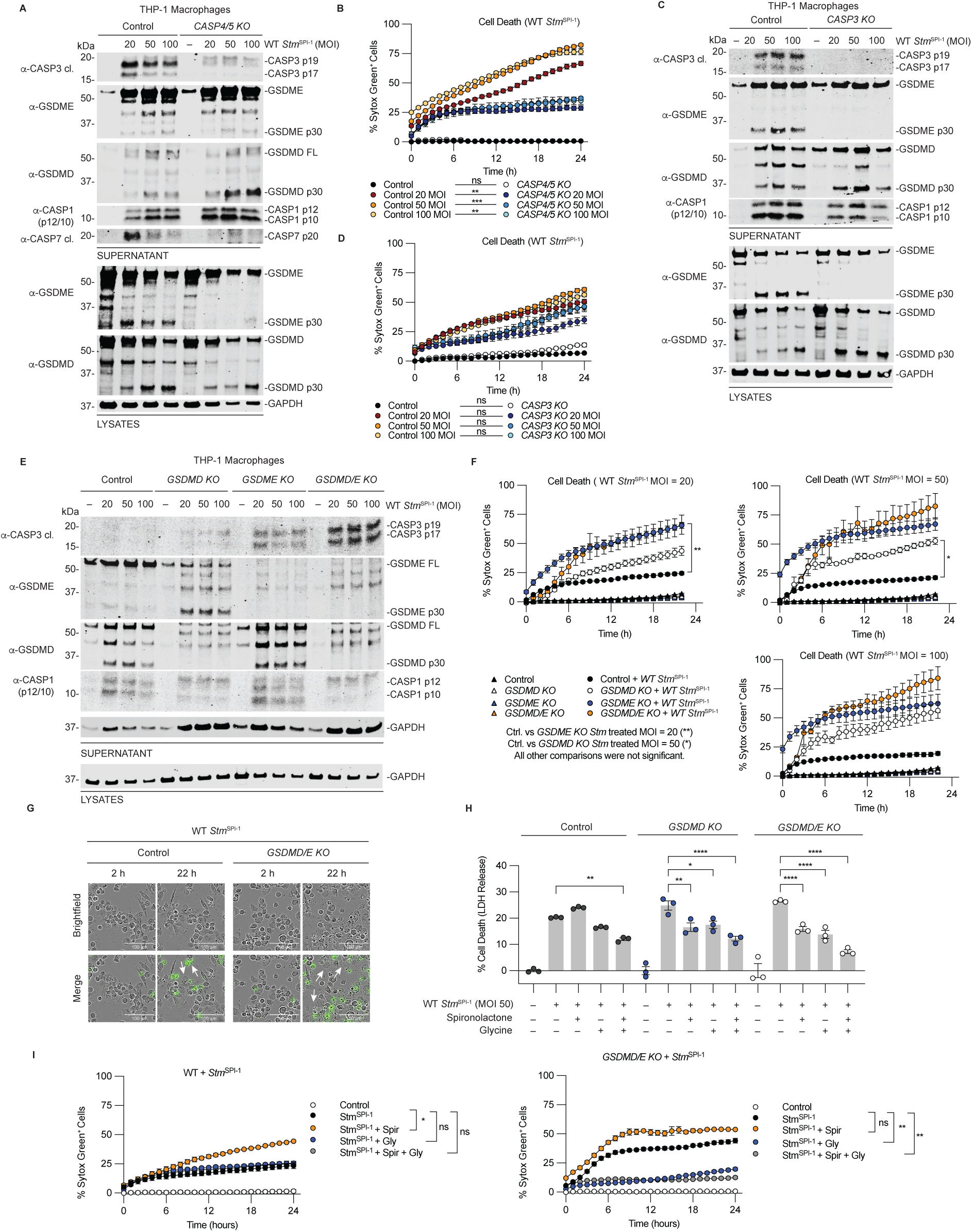
CASP3, and CASP4/5 are required for GSDME processing during non-canonical inflammasome activation by intracellular *Salmonella*. (A-D) WT and *CASP4/5* KO (A,B) or WT and *CASP3* KO THP-1 macrophages (C,D) were treated with the indicated MOI of WT SPI1-induced *Salmonella* for 24 h then supernatants and lysates were analyzed by immunoblotting (**A,C**), and cell death was measured by monitoring Sytox Green uptake (**B,D**). **(E-G)** WT, *GSDMD* KO, *GSDME* KO and *GSDMD/E* KO THP-1 macrophages were treated with the indicated MOI of WT SPI1-induced *Salmonella* for 24 h then supernatants and lysates were analyzed by immunoblotting (**E**), and cell death was measured by monitoring Sytox Green uptake (**F**). Examples of Images of WT and *GSDMD/E* KO cells at MOI 20 used to quantify Sytox Green uptake are shown in **G.** White arrows show pyroptotic morphology. **(H,I)**. WT, *GSDMD* KO, and *GSDMD/E* KO THP-1 macrophages were infected with WT SPI1-induced *Salmonella* at MOI = 50 in the presence or absences of pretreatment with the PANX1 (20 μM spironolactone) or NINJ1(50 mM glycine) inhibitors for 24 h then LDH release (H) and Sytox Green (**I**) were assessed. Data in H were analyzed by One-way Anova with Tukey’s multiple comparisons test. Data are means ± SEM of three independent replicates, and representative of at least three independent experiments.****P < 0.0001, ***P < 0.001, **P < 0.01, and *P < 0.05 by two-way ANOVA test with Tukey’s multiple comparison test comparing the control treated to KO treated samples at 24 h.

To determine the contribution of the gasdermin proteins to cell death, we infected WT, *GSDMD*, *GSDME*, or *GSDMD/E* KO THP-1 macrophages with *Stm*^SPI-1^ (Fig. 4E,F). We observed similar results in activation of CASP3/7 with *Salmonella* infection as with LPS transfections, but there were some notable differences in cell death (Fig. 4E,F). GSDME processing was increased in *GSDMD* KO cells and GSDMD processing was increased in *GSDME* KO cells unlike with LPS treatment where GSDMD processing was attenuated in *GSDME* KO cells, likely due to the activation of CASP1 by SPI-1-induced *Salmonella* (Fig. 4E). Indeed, CASP1 processing was similar in WT and *GSDME* KO cells, but was attenuated in *GSDMD* and *GSDMD/E* KO cells compared to WT cells. Strikingly, cell death, as assessed by Sytox Green uptake, was not attenuated in *GSDMD*, *GSDME*, or *GSDMD/E* KO cells at 24h post infection compared to WT cells and although not significant in all cases, trended higher (Fig. 4F).

Despite the loss of both pore-forming proteins, *GSDMD/E* KO cells displayed a lytic morphology consistent with gasdermin-independent rupture (Fig. 4G). This implies that enhanced CASP3/7 activity may drive rapid cell lysis, potentially through alternative pore-forming mechanisms such as Ninjurin-1 (NINJ1)-mediated membrane rupture^61,63^ or pannexin-1 (PANX1), a substrate of CASP3/7 that can form pores independently of GSDMD and GSDME^64^. To test this, WT, *GSDMD* KO and *GSDMD/E* KO cells were infected with *Stm*^SPI-1^ at MOI 50 after pretreatment with spironolactone, a pannexin-1 inhibitor, or glycine, an osmoprotectant that inhibits NINJ1-mediated lysis, or a combination of both^63,64^. Pretreatment with PANX1 or NINJ1 inhibitors separately had a modes impact but together, significantly reduced LDH release in all genotypes (Fig. 4H), supporting the notion that PANX1 channels and NINJ1-mediated rupture occurring downstream of apoptosis contribute to infection-induced lysis, like recently described^15,65^. However, inhibiting PANX1 and NINJ1 together had no impact on pore formation in WT cells, as assessed by Sytox Green uptake, but significantly inhibited Sytox Green uptake in *GSDMD/E* KO cells (Fig. 4I). PANX1 inhibition alone did not reduce Sytox Green uptake in *GSDMD/E* KO cells whereas NINJ1 inhibition did, hinting that lysis is mainly mediated by NINJ1. Collectively, these findings suggest that during intracellular LPS sensing, GSDMD is the dominant pore-forming protein mediating cell lysis, while both GSDMD and GSDME contribute to cytokine release. In contrast, during *Salmonella* infection, gasdermin-independent pathways can drive cell rupture downstream of non-canonical inflammasome mediated CASP3/7 activation.

### GSDMD and CASP3 are essential, but GSDME is dispensable for limiting *Salmonella* replication in human macrophages

Recent work revealed that CASP4 plays a role in limiting intracellular *Salmonella* replication, particularly at late stages of infection, though the mechanism remains unclear^15^. Activation of pyroptosis in response to infection is generally thought to be host-protective by limiting cellular replicative niches and activating cytokines that potentiate innate and adaptive immune responses. We hypothesized that CASP4 prevents replication by activating CASP3 to induce GSDME-mediated pyroptosis once bacterial burdens become high. To test the contribution of GSDME in preventing *Salmonella* replication, we infected WT, *GSDMD*, *GSDME,* and *GSDMD/E* KO THP-1 macrophages with GFP-expressing *Salmonella (GFP-Salmonella*) at increasing MOIs (MOI = 20, 50, or 100) and measured the change in GFP intensity during infection as a proxy for bacterial replication (Fig. 5). We collected images and quantified the fold change in *GFP-Salmonella* intensity at 24 h post infection relative to the start of imaging (t = 2 h) infection (Fig. 5A). Unexpectedly, GSDME was dispensable for limiting bacterial replication (Fig. 5B). Only *GSDMD* KO cells had significantly higher GFP-*Salmonella* burdens relative to WT infected cells (Fig. 5B), consistent with a recent study demonstrating that GSDMD is essential for limiting intracellular *Salmonella* replication^61^. *GSDME* KO cells had decreased burdens compared to WT cells, suggesting that GSDME may promote, instead of limit infection and replication. Consistent with this idea, *GSDMD/E* KO cells had similar bacterial burdens to WT cells and had decreased burdens compared to *GSDMD* KO cells. Cell death was assessed by LDH release and observed to be largely independent of both GSDMD and GSDME (Fig. 5C), consistent with contributions from PANX1 and NINJ1 activation.

**Figure 5.**
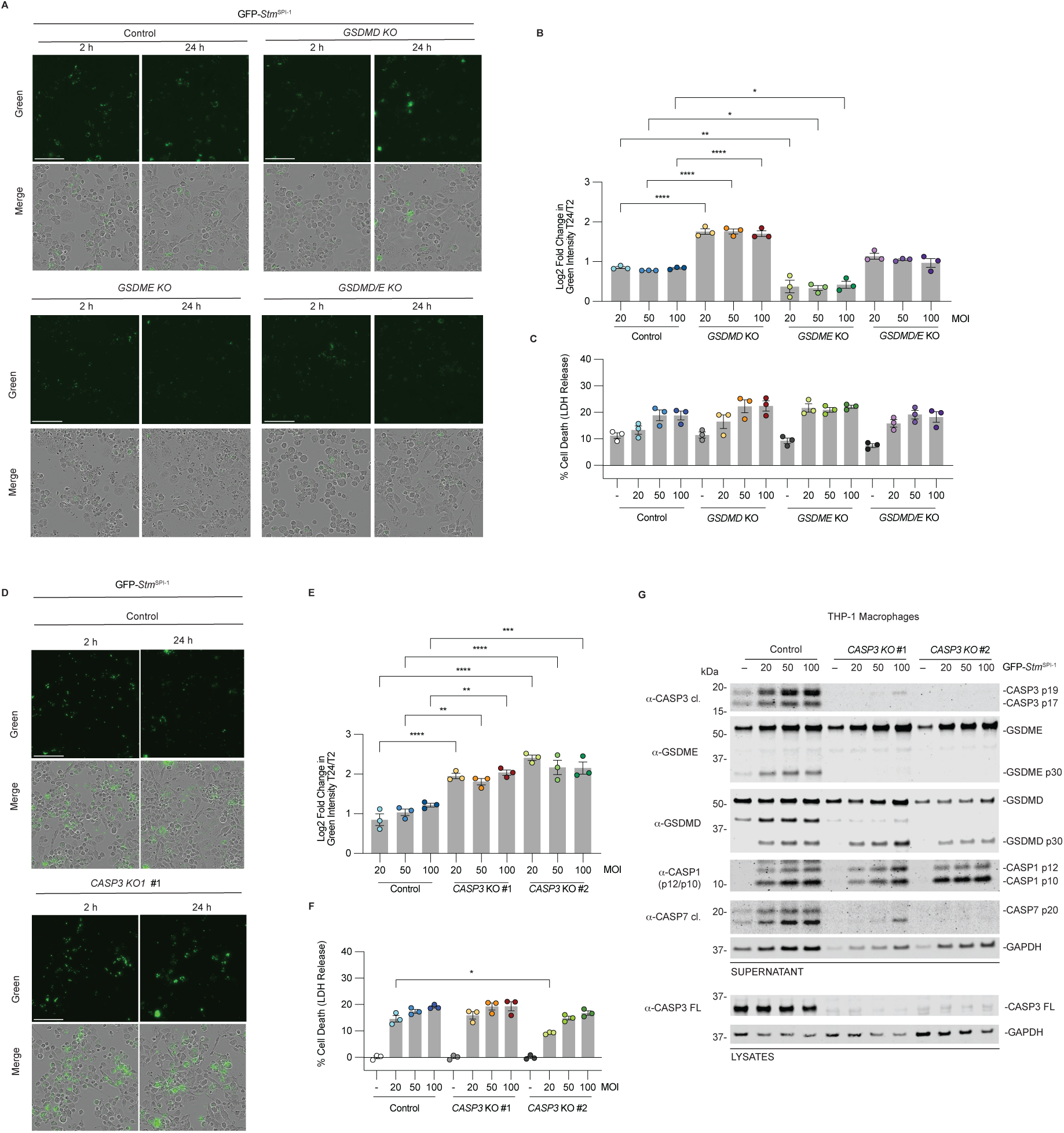
GSDMD and CASP3 are essential, but GSDME is dispensable for limiting *Salmonella* replication in human macrophages. (**A-C**) WT, *GSDMD* KO, *GSDME* KO and *GSDMD/E* KO THP-1 macrophages were treated with the indicated MOI of SPI-1-induced GFP-expressing *Salmonella*. Representative images of the GFP-*Salmonella* (MOI = 20) are shown in **A**. Quantification of the fold change in green intensity is depicted in **B**. After 24 h, LDH assays were performed **C**. (**D-F**) WT or *CASP3* KO THP-1 macrophages were treated with the indicated MOI of SPI1-induced GFP expressing *Salmonella*. Representative images of the GFP-*Salmonella* (MOI = 20) are shown in **D.** Quantification of the fold change in green intensity is depicted in **E**. After 24 h, LDH assays were performed **F**. Data are means ± SEM of three independent replicates, and representative of at least three independent experiments. ****P < 0.0001, ***P < 0.001, **P < 0.01, and *P < 0.05 by two-way ANOVA test with Tukey’s multiple comparison test comparing the control treated to KO treated samples at 24 h.

We next focused on the role of CASP3 in restricting bacterial replication, as *CASP7* KO cells had a confounding defect in GSDMD processing (Fig. 3G). WT and *CASP3* KO cells were infected with GFP-*Salmonella* and images of infected cells were collected to quantify the fold change in GFP intensity. In WT cells, GFP intensity remained stable, indicating replication control at all MOIs (Fig. 5D,E). In contrast, *CASP3* KO cells showed a significant increase in bacterial burden across all MOIs tested compared to WT cells (Fig. 5D,E). We tested two independent *CASP3* KO clones and LDH release at MOI = 20 was decreased in one clone compared to WT cells but unaffected at MOI = 50 and 100 (Fig. 5F). Together, these data indicate that in human macrophages, GSDMD and CASP3 are essential for restricting intracellular *Salmonella*, whereas GSDME is dispensable and may even facilitate replication.

### CASP3/7 and GSDME are cleaved in primary human macrophages during non-canonical inflammasome activation

To evaluate the physiological relevance of CASP4/5-induced CASP3/GSDME activation, we transfected primary human monocyte-derived macrophages (differentiated with macrophage colony stimulating factor (M-CSF)) with LPS ^66^. Consistent with observations in THP-1 cells, LPS transfection induced robust processing of CASP3/7 and GSDME in primary macrophages (Fig. 6A). Representative immunoblots from three independent donors are shown (Fig. 6A). Pretreatment with the NLRP3 inhibitor MCC950 blocked CASP1 activation and reduced GSDMD processing but did not alter CASP3/7 or GSDME cleavage, indicating that CASP3/7 activation is primarily mediated by CASP4/5, whereas GSDMD processing is largely downstream of CASP1 (Fig. 6A). Intracellular LPS-induced cell death and CASP3/7 activity were comparable between untreated and MCC950-treated cells (Fig. 6B-D), consistent with non-canonical inflammasome activation. Representative images from Donor 7 illustrate pyroptotic morphology and Sytox Green uptake used to monitor cell death kinetics (Fig. 6B). Aggregate analysis across all donors confirmed that cell death was unchanged by MCC950 treatment (Fig. 6E). Importantly, *Salmonella* infection induced CASP3/7, GSDME, and GSDMD processing in primary cells (Fig. 6F). Together, these findings demonstrate that intracellular LPS and *Salmonella* primarily engage CASP4/5 to induce GSDMD and CASP3/GSDME processing.

**Figure 6.**
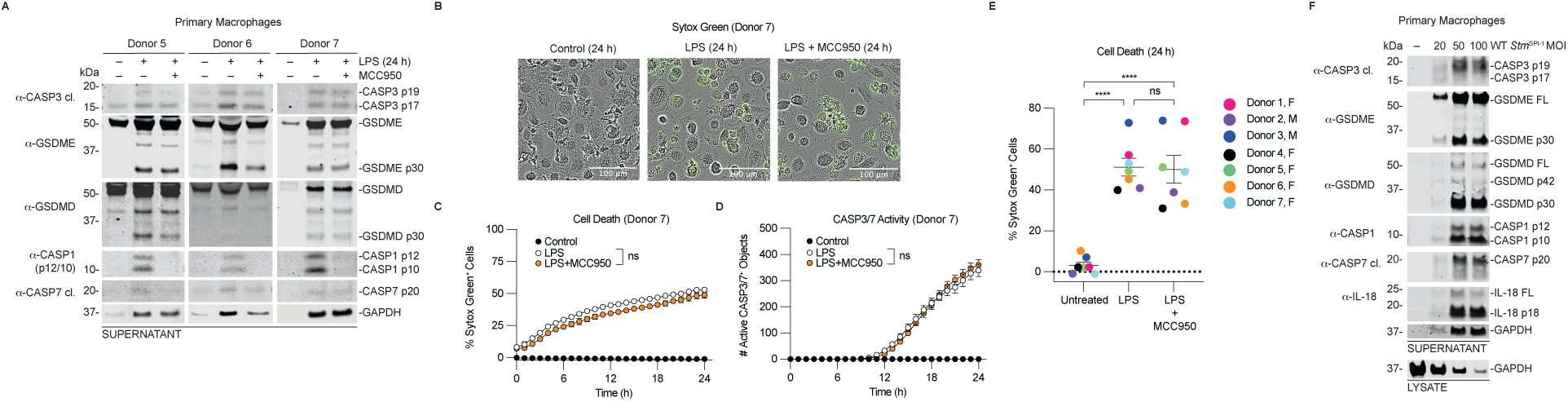
CASP3/7 and GSDME are cleaved in primary human macrophages during non-canonical inflammasome activation. (**A-E**) Primary monocytes were differentiated into macrophages and transfected with 25 μg/mL of LPS for 24 h. Some cells were pretreated with 10 μM MCC950 for 30 minutes before LPS transfection and supernatants were then analyzed by immunoblotting (**A**), cell death for a representative donor (Donor 7), as assessed by Sytox Green uptake is depicted in (**B**), and CASP3/7 activity CASP3/7 is depicted in (**C**). A summary of the Sytox Green uptake for all donors is shown in **E**. Data are means ± SEM of three or more independent experiments ****P < 0.0001, ***P < 0.001, **P < 0.01, and *P < 0.05 by two-way ANOVA test with Tukey’s multiple comparison test comparing the control vs treated samples at 24 h. Data in E were analyzed by One-way Anova with Tukey’s multiple comparisons test.

**Figure 7.**
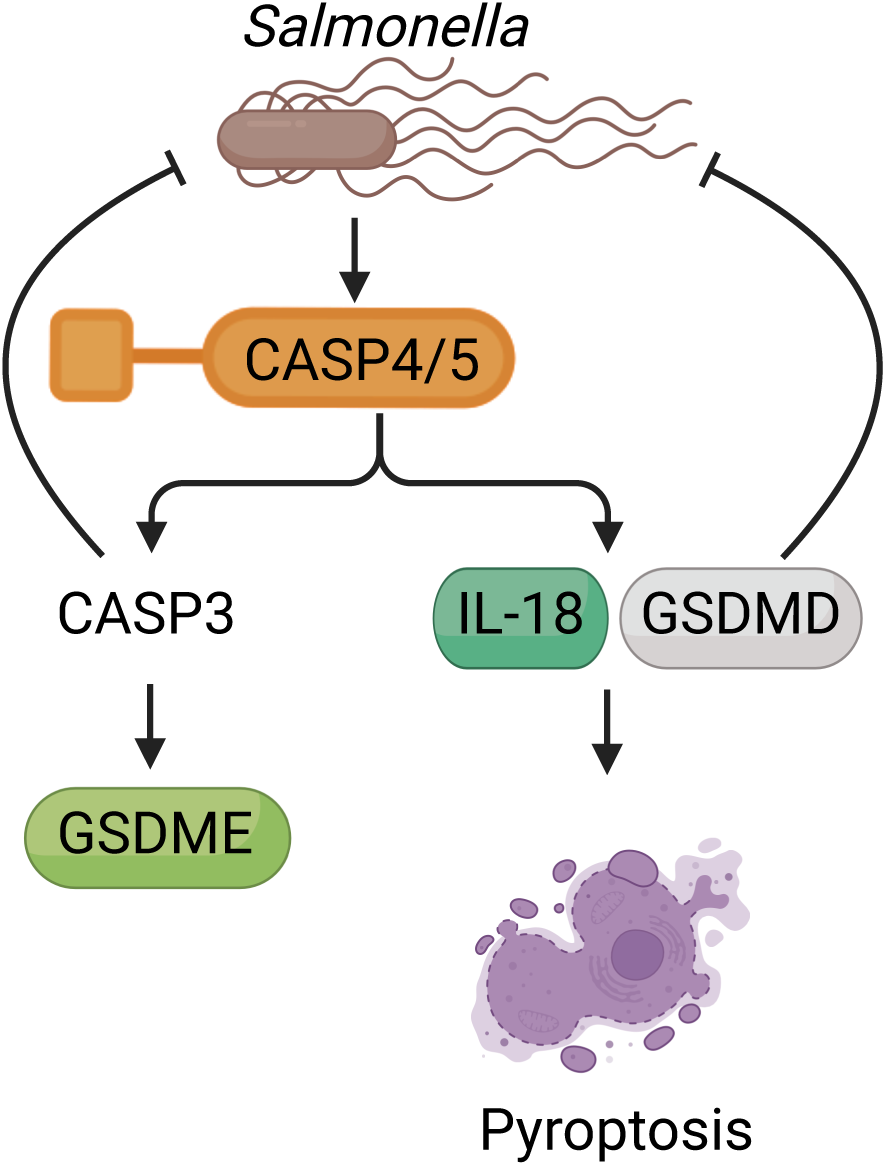
Proposed model of how non-canonical inflammasomes contribute to limiting intracellular *Salmonella* replication. Upon sensing LPS or *Salmonella*, CASP4/5 directly cleave CASP3/7 (CASP7 not shown), and CASP3 subsequently cleaves GSDME to activate it. However, GSDME does not appear to impact cell lysis, and is dispensable for controlling *Salmonella* replication. Notably, CASP3 activation is important for limiting intracellular *Salmonella* replication. CASP4/5 can also directly cleave GSDMD and IL-18 to induce pyroptosis, and GSDMD is essential for limiting bacterial replication. Figure generated in Biorender.

## Discussion

Cell death is central to maintaining cellular homeostasis and resolving infections, with inflammasome-mediated pyroptosis serving as a critical defense mechanism against intracellular pathogens^67^. Previous studies have revealed complex crosstalk between apoptotic and pyroptotic caspases, particularly in canonical inflammasome signaling, where CASP1 can indirectly or directly activate executioner CASP3 and -7 ^23,29,31,32,68-71^. These signaling cascades appears to be both species- and stimulus-dependent. Direct cleavage of CASP3/7 by CASP1 has been reported in human cells under certain conditions, such as dipeptidyl peptidase 8/9 inhibitor treatment^29^, while in mouse bone marrow–derived macrophages, CASP1-mediated CASP3/7 activation requires upstream CASP8 and CASP9 activation induced by CASP1-dependent Bid cleavage in response to inflammasome agonists like LPS plus nigericin (NLRP3 agonist), *Salmonella* (NLRC4 agonist), or poly(dA:dT) (AIM2 agonist) ^31,32^. Whether human non-canonical inflammasomes (CASP4/5) similarly activate apoptotic executioner caspases remained unclear.

In the present study, we demonstrate that human CASP4/5 directly cleave CASP3 and CASP7, and CASP3 subsequently cleaves GSDME, but our data suggests that most of the cell death is mediated by GSDMD. Unlike CASP1, CASP4/5 did not cleave Bid, and neither CASP8 nor CASP9 contributed to CASP3 activation during non-canonical inflammasome signaling. These data identify CASP4/5 as direct upstream activators of apoptotic executioner caspases in human cells. Although previous *in vitro* studies suggested that murine CASP11 (the ortholog of human CASP4) can cleave mouse CASP3/7, we did not observe significant CASP11 processing of human CASP3^70,72^. Other work indicates that in mouse macrophages, CASP11 triggers CASP1 activation, that then cleaves CASP3^71^. This points to species-specific differences in non-canonical inflammasome signaling, consistent with other observations that human CASP4/5, but not mouse CASP11, can cleave IL-18^20-22^.

Our data also reveal that pharmacological or genetic inhibition of CASP1 attenuated GSDMD processing while increasing GSDME cleavage, indicating that CASP4/5 preferentially drive CASP3/GSDME-mediated pore formation. Both GSDMD and GSDME contributed to IL-1β release as measured by immunoblotting, in agreement with prior observations^45^. However, IL-1β levels measured using an IL-1 receptor activation assay were not significantly reduced by single GSDMD or GSDME deletion, likely reflecting contributions from IL-1α release downstream of pore formation, which activates IL-1α via calpains^73^. Consistent with this, *GSDMD/E* double knockout cells transfected with LPS showed a reduction in released IL-1α/β compared to wild-type or single knockout cells, highlighting the collective role of these gasdermins in coordinating cytokine release.

The CASP4/5–CASP3/GSDME axis may function as a backup pathway when pathogens inhibit canonical CASP1-GSDMD signaling. For example, SARS-CoV-2 NSP5 protease cleaves GSDMD N-term to inactivate the pore forming ability, but the protease also activates CASP1 and CASP8, that subsequently activate CASP3 to induce GSDME-mediated pyroptosis^74^. Similarly, Enterovirus 3C protease cleaves and inactivates GSDMD N-term^75^, and gram-negative bacterial pathogen *Shigella flexneri* can degrade GSDMD, thereby impairing canonical pyroptosis and promoting infection^76^. Consistently, we observed increased CASP3/7 and GSDME cleavage in GSDMD-deficient cells, highlighting the compensatory nature of this pathway, similar to observations made by others^32,45^.

While GSDMD has long been recognized as the primary effector of LPS-induced pyroptosis via inflammatory caspases^17-19^, our data suggest that in human macrophages, most GSDMD processing is mediated by CASP1, which is activated downstream of the non-canonical pathway. In contrast, CASP4/5 preferentially induce GSDME-mediated pore formation via CASP3 activation, but the GSDME pores are sublytic. As CASP4/5 can cleave and inactivate IL-1ý^21^ and CASP3 is known to inactivate GSDMD^29^, IL-1β^77,78^, and IL-18^79^, it is possible that this pathway may have evolved to induce pyroptosis while limiting inflammation that could result in sepsis. GSDME-mediated pyroptosis may also have a distinct role in promoting a sublytic phase that enables rapid IL-1β release, which could have important ramifications for pathogen defense^45^.

Our data suggest reciprocal regulation between CASP1 and CASP3-induced pore formation. In CASP1-deficient cells, GSDME processing was increased, while GSDMD processing was increased in CASP3-deficient cells, suggesting that these pathways can compensate for one another to maintain overall cell lysis. This crosstalk may ensure robust inflammatory responses even when pathogens attempt to inhibit one arm of the pathway. Notably, GSDMD appears to play a dominant role in the kinetics of pyroptotic cell death, consistent with previous reports that GSDMD pore formation precedes apoptotic caspase activation during canonical inflammasome signaling^29,80^. However, this only applied in the context of when CASP4/5 were activated by LPS. In this study, neither GSDMD nor GSDME were required for *Salmonella* induced cell lysis at 24h post infection, contrasting with a recent study demonstrating that GSDMD and NINJ1 were essential for LDH release in THP-1 macrophages infected with *Salmonella^15^.* Differences in infection timepoint (6 h vs 24 h in this study), cell priming (e.g., Pam3CSK4 vs. unprimed in our study) and contributions of alternative pore-forming proteins such as NINJ1 or PANX1 may explain these discrepancies. Notably, recent work in mice found that LPS or *Escherichia coli* infection plus heat shock induced cell lysis that was independent of the typical pore-forming proteins such as, GSDMD, GSDME, and MLKL (mixed lineage kinase domain-like protein), but was critically dependent on NINJ1, which mediates plasma membrane rupture downstream of several cell death pathways^63,81^. Furthermore, this death only occurred with certain priming agents, such as poly(I:C) and LPS, but not Pam3CSK4^63^. Hence, it’s possible that under certain conditions, *Salmonella* can induce rapid cell lysis that is independent of GSDMD and GSDME. More studies using human cells are needed to understand the precise contributions of the different pore-forming proteins in pathogen defense.

Much of our understanding of the innate immune response to *Salmonella* has stemmed from work using mice, but emerging studies using human cells have begun to yield critical insights. Recent work in human macrophages demonstrated that CASP1-dependent pyroptosis is important for limiting intracellular *Salmonella* replication early during infection but CASP4 was important for restricting replication at later stages of infection^15^. Our data suggests that CASP4 inhibits replication in part by activating CASP3/7 to induce apoptosis and cell lysis that is likely dependent on NINJ1 or PANX1. Indeed, NINJ was reported to be important for cell death in THP1 macrophages infected with *Salmonella^15^*. Recent work also revealed that CASP3 can activate PANX1 to induce cell lysis independently of CASP1, CASP8, GSDMD, and GSDME^64^. Consistent with this notion, we found that *GSDMD/E* KO cells exhibited lytic morphology and GSDME was dispensable for inhibiting bacterial replication. Also in agreement, a recent report also noted that GSDME was dispensable, and that GSDMD is the only gasdermin that confers protection against acute *Salmonella* infection in the gut of mice^82^. Additional support for this mechanism stems from observations that apoptosis is critical for restricting *Legionella* infection and replication in dendritic cells^34^, and apoptosis was reported to be essential for limiting *Salmonella* replication in mice^65,83^. Collectively, this implies that apoptosis may play a broader role in preventing intracellular pathogen replication and future studies should focus on delineating the contributions of apoptosis and pyroptosis in restricting infection.

Controlled activation of pyroptosis protects the host from infection, and therefore it was surprising that GSDME was dispensable for protecting against *Salmonella* replication. If CASP3/GSDME-induced pyroptosis occurs without CASP1 activation, it could be detrimental to the host as it can facilitate bacterial egress and infection of bystander cells without the full immune response induced by CASP1 activation. In support of this idea, it was demonstrated that CASP11-mediated cell death in the absence of CASP1 activation was detrimental and resulted in increased susceptibility to *Salmonella* infection in mice^55^. Similarly, it was shown that GSDMD pores promote the release of the fungal pathogen *Candida albicans* from mouse macrophages, and inhibiting GSDMD was protective as it prevented *Candida albicans* escape while maintaining CASP1 activation and GSDMD-independent IL-1ý release^84^. Alternatively, if GSDME pores are sublytic, as has been suggested^45^, it could facilitate pathogen entry and render *GSDME* KO cells less susceptible to initial infection. We note that our GSDME KO cells were more resistant to infection than WT and *GSDMD* KO cells and required higher MOIs to observe similar levels of infections. Intriguingly, CASP3/GSDME-mediated pyroptosis appears to be critical for limiting viral replication^35^, suggesting that the impact on pathogen defense could be pathogen-specific. Importantly, we demonstrate that CASP3/GSDME are processed in primary human macrophages following non-canonical inflammasome activation, underscoring the physiological relevance of this pathway.

In conclusion, we identify CASP3/7 as direct substrates of human non-canonical inflammasomes (CASP4/5) in both immortalized and primary human macrophages. Mechanistically, CASP4/5 mediated cleavage of CASP3 initiates GSDME processing, but our data shows that CASP3 and GSDMD, but not GSDME, are critical for limiting intracellular *Salmonella* replication in THP-1 cells. Future work is needed to confirm the role of CASP4/5 and CASP3 on replication in primary cells. CASP3 activation can drive cell lysis that is independent of GSDMD and GSDME, likely through PANX and NINJ1, which have both been demonstrated to be important in limiting *Salmonella* replication^15,65^. Given that apoptosis has recently been reported to be important for infection control, our data suggests that this seemingly immune silent form of cell death may be more important than previously recognize in controlling infections. A key outstanding question is the molecular mechanism by which CASP4/5 recognize and cleave CASP3/7. Given that inflammatory caspases utilize exosites or tetrapeptide recognition sequences for substrate specificity^16^, it is likely that a combination of these factors underlies recognition. Indeed, recent complementary work using *in vitro* biochemical assays found that CASP3/7 are direct substrates of CASP4 and CASP5, and the primary mode of recognition and processing was dominated by the exosite interactions, offering critical insight into the molecular mechanism of recognition and processing^85^.

## Method Details

### Antibodies and Reagents

Antibodies used include: GSDMD Rabbit polyclonal Ab (Novus Biologicals, NBP2-33422), FLAG® M2 monoclonal Ab (Sigma, F3165), GAPDH Rabbit monoclonal Ab (Cell Signaling Tech, 14C10), CASP1 p12/10 Rabbit monoclonal Ab (Abcam, ab179515), CASP4 Rabbit polyclonal Ab (Cell Signaling Tech, 4450S), CASP5 Rabbit monoclonal Ab (Cell Signaling Tech, 46680S), Myc Mouse monoclonal Ab (Cell Signaling Tech, 2276S), CASP3 (Cell Signaling Tech 9664S), CASP3 (Cell Signaling Tech, 9662), CASP7 (Cell Signaling Tech, 8438S), CASP7 (Cell Signaling Tech, 12827S), GSDME (Abcam, ab215191), hIL-1β Goat Polyclonal Ab (R&D systems, AF-201-NA), hIL-18 Goat Ab (R&D systems, af2548), CASP8 (Cell Signaling Tech, 4790S), CASP9 (Cell Signaling Tech, 9508S), Bid (Cell Signaling Technology, 2002), IL-1ɑ (Novus, AF-200-NA), HA Rabbit monoclonal Ab (Cell Signaling Tech, 3724S), FKBP12 Rabbit polyclonal Ab (Abcam, ab24373). IRDye 800CW anti-rabbit (LICOR, 925-32211), IRDye 800CW anti-mouse (LI-COR, 925-32210), IRDye 680CW anti-rabbit (LI-COR, 925-68073), IRDye 680CW anti-mouse (LI-COR, 925-68072). Other reagents used include: LPS-EB Ultrapure (Invivogen, tlrl-3pelps), FuGENE HD (Promega, E2311), AP20187 (Tocris™ 6297/5).

### Recombinant Caspase Assays

All *in vitro* assays using recombinant caspases were performed in 20 μL reactions containing caspase assay buffer (20 mM PIPES, 100 mM NaCl, 10 mM DTT, 1mM EDTA, 0.1% CHAPS, 10% sucrose, pH 7.) as previously described^23^. 10 μg of plasmids encoding for GSDMD, CASP3, CASP7, or GSDME were transfected into HEK 293T cells (10 cm plates) using FuGENE HD Transfection Reagent (Promega, E2312) according to the manufacturer’s protocol. After 24 hours, cells were harvested and lysed by sonication for 10 second pulses for 60 seconds at 30% amplitude in PBS and centrifuged at 12,000 g to remove cell debris. Lysates were diluted 10x in caspase assay buffer and incubated with 0.25 activity units/μL of each indicated caspase. At the indicated time points, samples were mixed 1:1 with 2x Licor Protein Sample Loading Buffer (Neta Scientific, 928-40004) and boiled at 95 °C for 10 minutes, then analyzed by immunoblotting.

### Cell Culture

All cells were grown at 37 °C in a 5% CO2 atmosphere incubator. HEK 293T cells were purchased from ATCC and THP1 cells were obtained previously or acquired from the Bryant lab^21^ or Abbott lab^45^. HEK 293T cells were cultured in Dulbecco’s Modified Eagle’s Medium (DMEM) with L-glutamine and 10% fetal bovine serum (FBS). THP-1 cells were cultured in Roswell Park Memorial Institute (RPMI) medium 1640 with L-glutamine and 10% fetal bovine serum (FBS). Purified human monocytes from healthy donors were purchased from the University of Pennsylvania Human Immunology Core. Purified monocytes were cultured in RPMI + 10% FBS and 25 ng/ml rm-csf in 96 well plates at a density of 100,000 cells/well. Media was removed and replaced with Opti-MEM containing Sytox Green and MCC950 as indicated for 30 minutes prior to LPS transfection as previously described. Cell lines were negative for mycoplasma contamination.

### IncuCyte CASP3/7 activity and Sytox Green uptake assays

The kinetics of CASP3/7 activity and Sytox Green uptake were measured in cells using the IncuCyte S3 Live cell imaging system (Sartorius). THP-1 cells were resuspended in RPMI medium containing 50 ng/mL phorbol 12-myristate 12-acetate (PMA) and plated in black, clear-bottom 96-well plates at a density of 8 x 10^4^ cells/well. After 48 h, the media was then replaced with Opti-MEM (0.1 mL/well) supplemented with either the cell-permeable CASP3/7 green dye (final concentration: 10 µM (1:200 dilution)) or Sytox Green (final concentration: 0.2-0.5 µM), as indicated. A series of images were collected with a 10x objective at 1h intervals for 24 hours. The number of Sytox Green or CASP3/7 dye-positive cells in each image was determined using the IncuCyte S3 software with 3 different areas/well. Cell death was quantified relative to 0.1% Triton treated macrophages.

### Cloning

All plasmids were cloned using Gateway technology as previously described^21^. DNA encoding the indicated proteins were inserted between the *attR* recombination sites and shuttled into modified pLEX_307 vectors (Addgene) using Gateway technology (Thermo Fisher Scientific) according to the manufacturer’s instructions. Proteins expressed from these modified vectors contain an N-terminal *att*B1 linker (GSTSLYKKAGFAT) after any N-terminal tag or a C-terminal *att*B2 linker (DPAFLYKVVDI) preceding any C-terminal tag such as Myc or HA.

sgRNAs were designed using the Broad Institute’s web portal and cloned into the pXPR016_Hygro or plentiGuide-Puro vector (pXPR003) (Addgene #52963) as described previously^29,86^. The sgRNA sequences used were: h*CASP3 clone#1* 5′-CGTGGTACAGAACTGGACTG -3′ h*CASP3* clone#2 5′-TGTCGATGCAGCAAACCTCA -3′ h*CASP7* 5′-AGACAATCACGTCAAAACCC -3′ h*CASP9* 5′-ACATCGACTGTGAGAAGTTG -3′ Control *(GFP)* 5′-GGGCGAGGAGCTGTTCACCG-3′ All plasmids were verified by DNA sequencing (Genewiz).

### THP1 Knockout cell lines

Constructs were packaged into lentivirus in HEK 293T cells using the Fugene HD transfection reagent (Promega) and 2 μg of the vector, 2 μg psPAX2, and 0.2 μg pMD2-G. THP-1 were spinfected with virus for 2 h at 1000g at 30 °C supplemented 8 μg/mL polybrene. After 2 days, cells were selected for stable expression of S. pyogenes Cas9 (Addgene #52962) using blasticidin (5 μg/mL for THP-1) and for stable expression of sgRNAs using puromycin (0.5 μg/mL for THP-1) or hygromycin (100 μg/mL for THP-1). After 10 d, single cells were isolated by serial dilution and expanded. Cells were then harvested for immunoblotting or used in experiments.

### Generation of HEK 293T CASP3/7 double KO cells

HEK 293T cells were transiently transfected with plasmids encoding for CAS9 and sgRNAs targeting CASP3 (sgRNA Sequence: AATGGACTCTGGAATATCCC) and CASP7 (sgRNA Sequence: GCATCTATCCCCCCTAAAGT). After 48, the cells were selected with puromycin (1 μg/mL) until all control cells were no longer viable. Single cell clones were then obtained by serial dilution and were confirmed by immunoblotting.

### Transient Transfection

HEK 293T cells were seeded in 12-well culture plates at 0.25 × 10^6^ cells/well in DMEM. The following day, the indicated plasmids were mixed in Opti-MEM and transfected using FuGENE HD (Promega) according to the manufacturer’s protocol.

### LPS Transfection of human macrophages

THP-1 cells were resuspended in RPMI medium containing 50 ng/mL phorbol 12-myristate 12-acetate (PMA). Cells were plated in 96-well plates at a density of 8 x 10^4^ cells/well. After 48 h, the media was then replaced with Opti-MEM (0.1 mLs/well). Where indicated, cells were treated with MCC950 (10 µM) for 30 min at 37 °C 5% CO_2_ before LPS transfection. The LPS solution was prepared by adding LPS (25 mg/mL final concentration) and FuGENE (0.5% final concentration) to Opti-MEM. This solution was gently mixed by flicking and incubated for 30 min at room temperature before drop-wise addition to each well. The supernatants were collected separately, and cells were lysed by sonication 24 h post transfection. Harvested supernatants were precipitated by chloroform/methanol and analyzed by immunoblotting.

### HEK Blue IL-1α/β reporter assay

HEK-Blue IL-1β (Invivogen) reporter cells were used according to manufacturer’s protocols as previously described^21^ to quantify the amount of active IL-1α/β released. HEK Blue cells were seeded in 96 well plates at 5 x 10^4^ cells/well in Dulbecco’s Modified Eagle’s Medium (DMEM) with L-glutamine and 10% fetal bovine serum (FBS) to a total volume of 150 ml. 10 µL of supernatant from control or treated THP1 cells were added, and the samples were incubated overnight at 37°C in a 5% CO2 atmosphere incubator. A serial dilution of recombinant IL-1β (Invivogen) was included in all experiments to generate a standard curve which to calculate absolute levels of active IL-1β. 50 µL of supernatant was mixed with 150 µL of freshly prepared Quanti-Blue solution and incubated at room temperature for 10 minutes in the dark before reading absorbance at 620 nm using a Cytation 5 plate reader (Bio Tek).

### *Salmonella* Infection of human macrophages

*Salmonella* Typhimurium SL1344 or SL1344 pDiGc^87^ was streaked to isolation from frozen stock on LB + Streptomycin (100µg/ml) agar plates. Carbenicillin (100µg/ml) was included for all studies with SL1344 pDiGc. For SPI-I induced bacteria, single colonies were inoculated in LB + Streptomycin broth overnight (∼18h). The overnight culture was back diluted 40-fold into LB + Streptomycin broth containing 300 mM NaCl. After three hours bacteria were washed 2x in sterile PBS and cell density was determined by quantifying OD_600_. MOI was prepared according to OD and bacterial cells were resuspended in PBS. 10µL suspension of bacteria was added to macrophages in triplicate in 96-well plates. Macrophages were spun for 10 min at 400g to allow infection and then placed in incubator for 1h. Media was then removed and replaced with fresh RPMI without FBS containing gentamicin (100 μg/ml). After 1h, media was removed and wells were washed 3x with PBS and replaced with fresh RPMI without FBS containing gentamicin (10 μg/ml). GFP fluorescence was monitored using IncuCyte live cell imaging for 24h.

For SL1344 (non-fluorescent) infection, after 1h 100µg/ml gentamicin, media was replaced with fresh RPMI without FBS containing gentamicin (10 μg/ml) and Sytox Green (0.5 µM). Cell death was quantified relative to 0.1% Triton treated macrophages.

For stationary phase bacterial infections, single colonies were inoculated in LB + Streptomycin for 2 days. MOI were prepared as described above and 10µL suspension of bacteria were added to macrophages in triplicate in 96 well plates. Macrophages were spun for 10 min at 400g to allow infection and then placed in incubator for 2h.

### LDH Assays

Supernatants were harvested for LDH analyses at 24 h after bacterial infections and analyzed using the Cyquant LDH Cytotoxicity Assay (Thermo Scientific) according to the manufacturer’s protocol. LDH activity was quantified relative to a lysis control where cells were lysed using Triton-X at the beginning of each experiment.

### Western Blotting

Protein samples were run on Nupage 4-12% Bis-Tris Midi Protein Gel (Thermo Scientific, WG1403BOX) at 175 volts. Proteins were transferred to 0.45 µm nitrocellulose membranes (BioRad,1704271) at 25 volts for 7 minutes on the Biorad Transblot Turbo System. Membranes were blocked using LICOR Intercept blocking buffer (Neta scientific, 927-70010) for 1 hour and stained with primary antibodies at 1:1000 concentration in 50% Licor blocking buffer and 50% TBS buffer with 0.1% Tween for 1 hour at room temperature or overnight at 4C. Membranes were washed 3x 15 minutes, followed by incubation with Donkey Anti-Mouse/Rabbit/Goat IgG Polyclonal Antibody (IRDye® 800CW) for 1 hour at room temperature. Membranes were then washed 3x10 minutes and imaged on an Odyssey M Imaging System (LI-COR Biosciences). Images were analyzed with Empiria studio version 3.2 (Licor), and brightness, contrast, and tone parameters were adjusted uniformly for entire membranes on Adobe Photoshop.

### Statistical Analysis

Data and statistical analysis were performed using GraphPad Prism 10 software. Statistical analysis was determined using One-way or Two-way ANOVA with multiple comparisons test. Further details are listed under each figure legend.

## Author contributions CRediT

Conceptualization: CYT, Ma.Ku., CMB Methodology: CYT, Ma.Ku, CMB, Investigation: Ma.Ku., CMB, ABM, PME, CM, SC, Ma.Ka., Mi.Ka., WY, TJW, RCP, BMD Visualization: CYT, Ma.Ku., CMB, Funding acquisition: CYT, CMB, Project administration: CYT Data curation: CYT, Ma.Ku., CMB, Formal Analysis: CYT, Ma.Ku., CMB, ABM, PME, CM, SC, Ma.Ka., Mi.Ka., Resources: CYT Supervision: CYT, Writing – original draft: CYT, Ma.Ku., CMB, Writing – review & editing: CYT, Ma.Ku., CMB.

## Acknowledgements

We thank Dr. Derek Abbott for providing *GSDMD* KO, *GSDME* KO, *GSDMD/E* KO, and *CASP8* KO THP-1 cells. We thank Dr. Igor Brodsky for providing strain SL1344 pDiGc for infection studies. The authors thank Emily Cento, Zhilin Chen, Max A. Eldabbas, and Emileigh Maddox of the Human Immunology Core and the Division of Transfusion Medicine and Therapeutic Pathology at the Perelman School of Medicine at the University of Pennsylvania for providing de-identified monocytes that were purified from healthy donor apheresis using StemCell RosetteSep™ kits. The HIC is supported in part by NIH P30 AI045008 and P30 CA016520. HIC RRID: SCR_022380. This work was supported by an NIGMS Maximizing Investigator’s Research Award (MIRA) Grant# 1R35GM155239-01 (CYT), and the Colton Center for Autoimmunity at Penn (CYT). CMB is a Penn Provost Postdoctoral Fellow and is supported by Burroughs Wellcome Fund (Grant# 1054907).

## Conflicts of Interest

The authors declare no conflicts of interests.

## Notes

### Competing Interest Statement

The authors have declared no competing interest.

### Summary of Updates

This version of the manuscript has been revised to update the model in Figure 7.

## References

1. Fleischmann-Struzek, C., and Rudd, K. (2023). Challenges of assessing the burden of sepsis. Med Klin Intensivmed Notfmed 118, 68–74. 10.1007/s00063-023-01088-7.

2. Rudd, K.E., Johnson, S.C., Agesa, K.M., Shackelford, K.A., Tsoi, D., Kievlan, D.R., Colombara, D.V., Ikuta, K.S., Kissoon, N., Finfer, S., et al. (2020). Global, regional, and national sepsis incidence and mortality, 1990-2017: analysis for the Global Burden of Disease Study. Lancet 395, 200–211. 10.1016/S0140-6736(19)32989-7.

3. Martinon, F., Burns, K., and Tschopp, J. (2002). The inflammasome: a molecular platform triggering activation of inflammatory caspases and processing of proIL-beta. Mol. Cell 10, 417–426. 10.1016/s1097-2765(02)00599-3.

4. Broz, P., and Dixit, V.M. (2016). Inflammasomes: Mechanism of assembly, regulation and signalling. Nature Reviews Immunology 16, 407–420. 10.1038/nri.2016.58.

5. Barnett, K.C., Li, S., Liang, K., and Ting, J.P.Y. (2023). A 360° view of the inflammasome: Mechanisms of activation, cell death, and diseases. Cell 186, 2288–2312. 10.1016/j.cell.2023.04.025.

6. Shi, J., Zhao, Y., Wang, Y., Gao, W., Ding, J., Li, P., Hu, L., and Shao, F. (2014). Inflammatory caspases are innate immune receptors for intracellular LPS. Nature 514, 187–192. 10.1038/nature13683.

7. Hagar, J.A., Powell, D.A., Aachoui, Y., Ernst, R.K., and Miao, E.A. (2013). Cytoplasmic LPS Activates Caspase-11: Implications in TLR4-Independent Endotoxic Shock. Science 341, 1250–1253. 10.1126/science.1240988.

8. Rühl, S., and Broz, P. (2015). Caspase-11 activates a canonical NLRP3 inflammasome by promoting K+ efflux. European Journal of Immunology 45, 2927–2936. 10.1002/eji.201545772.

9. Muñoz-Planillo, R., Kuffa, P., Martínez-Colón, G., Smith, B., Rajendiran, T., and Núñez, G. (2013). K+ Efflux Is the Common Trigger of NLRP3 Inflammasome Activation by Bacterial Toxins and Particulate Matter. Immunity 38, 1142–1153. 10.1016/j.immuni.2013.05.016.

10. Taabazuing, C.Y., Griswold, A.R., and Bachovchin, D.A. (2020). The NLRP1 and CARD8 inflammasomes. Immunological Reviews 297, 13–25. 10.1111/imr.12884.

11. Kulsuptrakul, J., Turcotte, E.A., Emerman, M., and Mitchell, P.S. (2023). A human-specific motif facilitates CARD8 inflammasome activation after HIV-1 infection. eLife 12, e84108. 10.7554/eLife.84108.

12. Majowicz, S.E., Musto, J., Scallan, E., Angulo, F.J., Kirk, M., O’Brien, S.J., Jones, T.F., Fazil, A., Hoekstra, R.M., and Studies, f.t.I.C.o.E.D.B.o.I. (2010). The Global Burden of Nontyphoidal Salmonella Gastroenteritis. Clinical Infectious Diseases 50, 882–889. 10.1086/650733.

13. Rathinam, V.A.K., Zhao, Y., and Shao, F. (2019). Innate immunity to intracellular LPS. Nature Immunology 20, 527–533. 10.1038/s41590-019-0368-3.

14. Casson, C.N., Yu, J., Reyes, V.M., Taschuk, F.O., Yadav, A., Copenhaver, A.M., Nguyen, H.T., Collman, R.G., and Shin, S. (2015). Human caspase-4 mediates noncanonical inflammasome activation against gram-negative bacterial pathogens. Proceedings of the National Academy of Sciences of the United States of America 112, 6688–6693. 10.1073/pnas.1421699112.

15. Egan, M.S., O’Rourke, E.A., Mageswaran, S.K., Zuo, B., Martynyuk, I., Demissie, T., Hunter, E.N., Bass, A.R., Chang, Y.-W., Brodsky, I.E., and Shin, S. (2024). Inflammasomes primarily restrict cytosolic Salmonella replication within human macrophages. eLife.

16. Exconde, P.M., Bourne, C.M., Kulkarni, M., Discher, B.M., and Taabazuing, C.Y. (2024). Inflammatory caspase substrate specificities. mBio 15, e0297523. 10.1128/mbio.02975-23.

17. Kayagaki, N., Stowe, I.B., Lee, B.L., O’Rourke, K., Anderson, K., Warming, S., Cuellar, T., Haley, B., Roose-Girma, M., Phung, Q.T., et al. (2015). Caspase-11 cleaves gasdermin D for non-canonical inflammasome signalling. Nature 526, 666–671. 10.1038/nature15541.

18. He, W.T., Wan, H., Hu, L., Chen, P., Wang, X., Huang, Z., Yang, Z.H., Zhong, C.Q., and Han, J. (2015). Gasdermin D is an executor of pyroptosis and required for interleukin-1β secretion. Cell Research 25, 1285–1298. 10.1038/cr.2015.139.

19. Shi, J., Zhao, Y., Wang, K., Shi, X., Wang, Y., Huang, H., Zhuang, Y., Cai, T., Wang, F., and Shao, F. (2015). Cleavage of GSDMD by inflammatory caspases determines pyroptotic cell death. Nature 526, 660–665. 10.1038/nature15514.

20. Devant, P., Dong, Y., Mintseris, J., Ma, W., Gygi, S.P., Wu, H., and Kagan, J.C. (2023). Structural insights into cytokine cleavage by inflammatory caspase-4. Nature 624, 451–459. 10.1038/s41586-023-06751-9.

21. Exconde, P.M., Hernandez-Chavez, C., Bourne, C.M., Richards, R.M., Bray, M.B., Lopez, J.L., Srivastava, T., Egan, M.S., Zhang, J., Yoo, W., et al. (2023). The tetrapeptide sequence of IL-18 and IL-1β regulates their recruitment and activation by inflammatory caspases. Cell Reports 42, 113581.

22. Shi, X., Sun, Q., Hou, Y., Zeng, H., Cao, Y., Dong, M., Ding, J., and Shao, F. (2023). Recognition and maturation of IL-18 by caspase-4 noncanonical inflammasome. Nature 624, 442–450. 10.1038/s41586-023-06742-w.

23. Bourne, C.M., Raniszewski, N.R., Mahale, A.B., Kulkarni, M., Exconde, P.M., Liu, S., Yost, W., Wrong, T.J., Patio, R.C., Kardhashi, M., et al. (2025). A Potent Inhibitor of Caspase-8 Based on the IL-18 Tetrapeptide Sequence Reveals Shared Specificities between Inflammatory and Apoptotic Initiator Caspases. ACS Bio & Med Chem Au. 10.1021/acsbiomedchemau.4c00146.

24. Miao, E.A., Leaf, I.A., Treuting, P.M., Mao, D.P., Dors, M., Sarkar, A., Warren, S.E., Wewers, M.D., and Aderem, A. (2010). Caspase-1-induced pyroptosis is an innate immune effector mechanism against intracellular bacteria. Nat Immunol 11, 1136–1142. 10.1038/ni.1960.

25. Ewald, S.E., Chavarria-Smith, J., and Boothroyd, J.C. (2014). NLRP1 is an inflammasome sensor for Toxoplasma gondii. Infection and Immunity 82, 460–468. 10.1128/IAI.01170-13.

26. Terra, J.K., France, B., Cote, C.K., Jenkins, A.L., Bozue, J.A., Welkos, S.L., Bhargava, R., Ho, C.L., Mehrabian, M., Pan, C., et al. (2011). Cutting Edge: Resistance to Bacillus anthracis Infection Mediated by a Lethal Toxin Sensitive Allele of Nalp1b/Nlrp1b. PLoS Pathogens 7, 17–20. 10.1371/journal.ppat.1002469.

27. Jorgensen, I., Zhang, Y., Krantz, B.A., and Miao, E.A. (2016). Pyroptosis triggers pore-induced intracellular traps (PITs) that capture bacteria and lead to their clearance by efferocytosis. The Journal of Experimental Medicine 213, 2113–2128. 10.1084/jem.20151613.

28. Miao, E.A., Leaf, I.A., Treuting, P.M., Mao, D.P., Dors, M., Sarkar, A., Warren, S.E., Wewers, M.D., and Aderem, A. (2010). Caspase-1-induced pyroptosis is an innate immune effector mechanism against intracellular bacteria. Nature Immunology 11, 1136–1142. 10.1038/ni.1960.

29. Taabazuing, C.Y., Okondo, M.C., and Bachovchin, D.A. (2017). Pyroptosis and Apoptosis Pathways Engage in Bidirectional Crosstalk in Monocytes and Macrophages. Cell Chemical Biology 24, 507–514. 10.1016/j.chembiol.2017.03.009.

30. Lamkanfi, M., Kanneganti, T.-D., Van Damme, P., Vanden Berghe, T., Vanoverberghe, I., Vandekerckhove, J., Vandenabeele, P., Gevaert, K., and Núñez, G. (2008). Targeted peptidecentric proteomics reveals caspase-7 as a substrate of the caspase-1 inflammasomes. Molecular & cellular proteomics : MCP 7, 2350–2363. 10.1074/mcp.M800132-MCP200.

31. Tsuchiya, K., Nakajima, S., Hosojima, S., Thi Nguyen, D., Hattori, T., Manh Le, T., Hori, O., Mahib, M.R., Yamaguchi, Y., Miura, M., et al. (2019). Caspase-1 initiates apoptosis in the absence of gasdermin D. Nature Communications 10. 10.1038/s41467-019-09753-2.

32. Heilig, R., Dilucca, M., Boucher, D., Chen, K.W., Hancz, D., Demarco, B., Shkarina, K., and Broz, P. (2020). Caspase-1 cleaves Bid to release mitochondrial SMAC and drive secondary necrosis in the absence of GSDMD. Life science alliance 3, e202000735. 10.26508/lsa.202000735.

33. Nogueira, C.V., Lindsten, T., Jamieson, A.M., Case, C.L., Shin, S., Thompson, C.B., and Roy, C.R. (2009). Rapid pathogen-induced apoptosis: a mechanism used by dendritic cells to limit intracellular replication of Legionella pneumophila. PLoS Pathog 5, e1000478. 10.1371/journal.ppat.1000478.

34. Vázquez Marrero Víctor, R., Doerner, J., Wodzanowski Kimberly, A., Zhang, J., Lu, A., Boyer Frankie, D., Vargas, I., Hossain, S., Kammann Karly, B., Dresler Madison, V., and Shin, S. (2025). Dendritic cells activate pyroptosis and effector-triggered apoptosis to restrict Legionella infection. mBio 16, e01257–01225. 10.1128/mbio.01257-25.

35. Orzalli, M.H., Prochera, A., Payne, L., Smith, A., Garlick, J.A., and Kagan, J.C. (2021). Virus-mediated inactivation of anti-apoptotic Bcl-2 family members promotes Gasdermin-E-dependent pyroptosis in barrier epithelial cells. Immunity 54, 1447–1462 e1445. 10.1016/j.immuni.2021.04.012.

36. Kovacs, S.B., and Miao, E.A. (2017). Gasdermins: Effectors of Pyroptosis. Trends Cell Biol 27, 673–684. 10.1016/j.tcb.2017.05.005.

37. Davies, C.W., Stowe, I., Phung, Q.T., Ho, H., Bakalarski, C.E., Gupta, A., Zhang, Y., Lill, J.R., Payandeh, J., Kayagaki, N., and Koerber, J.T. (2021). Discovery of a caspase cleavage motif antibody reveals insights into noncanonical inflammasome function. Proceedings of the National Academy of Sciences - PNAS 118, 1. 10.1073/pnas.2018024118.

38. Wang, Y., Gao, W., Shi, X., Ding, J., Liu, W., He, H., Wang, K., and Shao, F. (2017). Chemotherapy drugs induce pyroptosis through caspase-3 cleavage of a gasdermin. Nature 547, 99–103. 10.1038/nature22393.

39. Green, D.R. (2022). Caspases and Their Substrates. Cold Spring Harbor perspectives in biology 14, a041012. 10.1101/cshperspect.a041012.

40. Rano, T.A., Timkey, T., Peterson, E.P., Rotonda, J., Nicholson, D.W., Becker, J.W., Chapman, K.T., and Thornberry, N.A. (1997). A combinatorial approach for determining protease specificities: application to interleukin-1β converting enzyme (ICE). Chemistry & biology 4, 149–155. 10.1016/S1074-5521(97)90258-1.

41. Talanian, R.V., Quinlan, C., Trautz, S., Hackett, M.C., Mankovich, J.A., Banach, D., Ghayur, T., Brady, K.D., and Wong, W.W. (1997). Substrate Specificities of Caspase Family Proteases. The Journal of biological chemistry 272, 9677–9682. 10.1074/jbc.272.15.9677.

42. Ponder, K.G., and Boise, L.H. (2019). The prodomain of caspase-3 regulates its own removal and caspase activation. Cell Death Discov 5, 56. 10.1038/s41420-019-0142-1.

43. Rogers, C., Fernandes-Alnemri, T., Mayes, L., Alnemri, D., Cingolani, G., and Alnemri, E.S. (2017). Cleavage of DFNA5 by caspase-3 during apoptosis mediates progression to secondary necrotic/pyroptotic cell death. Nat Commun 8, 14128. 10.1038/ncomms14128.

44. Baker, P.J., Boucher, D., Bierschenk, D., Tebartz, C., Whitney, P.G., D’Silva, D.B., Tanzer, M.C., Monteleone, M., Robertson, A.A., Cooper, M.A., et al. (2015). NLRP3 inflammasome activation downstream of cytoplasmic LPS recognition by both caspase-4 and caspase-5. Eur J Immunol 45, 2918–2926. 10.1002/eji.201545655.

45. Zhou, B., and Abbott, D.W. (2021). Gasdermin E permits interleukin-1 beta release in distinct sublytic and pyroptotic phases. Cell Rep 35, 108998. 10.1016/j.celrep.2021.108998.

46. Coll, R.C., Hill, J.R., Day, C.J., Zamoshnikova, A., Boucher, D., Massey, N.L., Chitty, J.L., Fraser, J.A., Jennings, M.P., Robertson, A.A.B., and Schroder, K. (2019). MCC950 directly targets the NLRP3 ATP-hydrolysis motif for inflammasome inhibition. Nat Chem Biol 15, 556–559. 10.1038/s41589-019-0277-7.

47. Julien, O., and Wells, J.A. (2017). Caspases and their substrates. Cell death and differentiation 24, 1380–1389. 10.1038/cdd.2017.44.

48. Singh, R., Letai, A., and Sarosiek, K. (2019). Regulation of apoptosis in health and disease: the balancing act of BCL-2 family proteins. Nat Rev Mol Cell Biol 20, 175–193. 10.1038/s41580-018-0089-8.

49. Billen, L.P., Shamas-Din, A., and Andrews, D.W. (2008). Bid: a Bax-like BH3 protein. Oncogene 27 *Suppl 1*, S93–104. 10.1038/onc.2009.47.

50. Esposti, M., Ferry, G., Masdehors, P., Boutin, J., Hickman, J., and Dive, C. (2003). Post-translational Modification of Bid Has Differential Effects on Its Susceptibility to Cleavage by Caspase 8 or Caspase 3. The Journal of biological chemistry 278, 15749–15757. 10.1074/jbc.M209208200.

51. Orning, P., Lien, E., and Fitzgerald, K.A. (2019). Gasdermins and their role in immunity and inflammation. The Journal of Experimental Medicine *In Press*, jem.20190545-jem.20190545. 10.1084/jem.20190545.

52. Li, S., Bracey, S., Liu, Z., and Xiao, T.S. (2023). Regulation of gasdermins in pyroptosis and cytokine release. Adv Immunol 158, 75–106. 10.1016/bs.ai.2023.03.002.

53. Zhou, Z., He, H., Wang, K., Shi, X., Wang, Y., Su, Y., Wang, Y., Li, D., Liu, W., Zhang, Y., et al. (2020). Granzyme A from cytotoxic lymphocytes cleaves GSDMB to trigger pyroptosis in target cells. Science 368. 10.1126/science.aaz7548.

54. Zhang, Z., Zhang, Y., Xia, S., Kong, Q., Li, S., Liu, X., Junqueira, C., Meza-Sosa, K.F., Mok, T.M.Y., Ansara, J., et al. (2020). Gasdermin E suppresses tumour growth by activating anti-tumour immunity. Nature 579, 415–420. 10.1038/s41586-020-2071-9.

55. Broz, P., Ruby, T., Belhocine, K., Bouley, D.M., Kayagaki, N., Dixit, V.M., and Monack, D.M. (2012). Caspase-11 increases susceptibility to Salmonella infection in the absence of caspase-1. Nature 490, 288–291. 10.1038/nature11419.

56. Miao, E.A., Mao, D.P., Yudkovsky, N., Bonneau, R., Lorang, C.G., Warren, S.E., Leaf, I.A., and Aderem, A. (2010). Innate immune detection of the type III secretion apparatus through the NLRC4 inflammasome. Proc Natl Acad Sci U S A 107, 3076–3080. 10.1073/pnas.0913087107.

57. Broz, P., Ohlson, M.B., and Monack, D.M. (2012). Innate immune response to Salmonella typhimurium, a model enteric pathogen. Gut Microbes 3, 62–70. 10.4161/gmic.19141.

58. Broz, P., Newton, K., Lamkanfi, M., Mariathasan, S., Dixit, V.M., and Monack, D.M. (2010). Redundant roles for inflammasome receptors NLRP3 and NLRC4 in host defense against Salmonella. J Exp Med 207, 1745–1755. 10.1084/jem.20100257.

59. Naseer, N., Egan, M.S., Reyes Ruiz, V.M., Scott, W.P., Hunter, E.N., Demissie, T., Rauch, I., Brodsky, I.E., and Shin, S. (2022). Human NAIP/NLRC4 and NLRP3 inflammasomes detect Salmonella type III secretion system activities to restrict intracellular bacterial replication. PLoS Pathogens 18. 10.1371/journal.ppat.1009718.

60. Naseer, N., Zhang, J., Bauer, R., Constant, D.A., Nice, T.J., Brodsky, I.E., Rauch, I., and Shin, S. (2022). Salmonella enterica Serovar Typhimurium Induces NAIP/NLRC4-And NLRP3/ASC-Independent, Caspase-4-Dependent Inflammasome Activation in Human Intestinal Epithelial Cells. Infection and Immunity 90. 10.1128/iai.00663-21.

61. Egan, M.S., O’Rourke, E.A., Mageswaran, S.K., Zuo, B., Martynyuk, I., Demissie, T., Hunter, E.N., Bass, A.R., Chang, Y.-W., Brodsky, I.E., and Shin, S. (2023). Inflammasomes primarily restrict cytosolic Salmonella replication within human macrophages. eLife 12, 90107.90101. 10.7554/elife.90107.1.

62. Lou, L., Zhang, P., Piao, R., and Wang, Y. (2019). Salmonella Pathogenicity Island 1 (SPI-1) and Its Complex Regulatory Network. Front Cell Infect Microbiol 9, 270. 10.3389/fcimb.2019.00270.

63. Han, J.H., Karki, R., Malireddi, R.K.S., Mall, R., Sarkar, R., Sharma, B.R., Klein, J., Berns, H., Pisharath, H., Pruett-Miller, S.M., et al. (2024). NINJ1 mediates inflammatory cell death, PANoptosis, and lethality during infection conditions and heat stress. Nat Commun 15, 1739. 10.1038/s41467-024-45466-x.

64. Zhou, B., Ryder, C.B., Dubyak, G.R., and Abbott, D.W. (2022). Gasdermins and pannexin-1 mediate pathways of chemotherapy-induced cell lysis in hematopoietic malignancies. Science Signaling 15, eabl6781. doi:10.1126/scisignal.abl6781.

65. Herrmann, B.I., Zamba-Campero, M., Sillas, R.G., Yost, W.W., Peterson, L.W., Roncaioili, J.L., Rankin, S.C., Schifferli, D.M., Ravichandran, K., and Brodsky, I.E. (2025). Innate detection of Salmonella replication triggers caspase-8-dependent apoptosis via TLR-driven TNF signaling and NLRC4-mediated sensing of the SPI-2 Type III secretion system. bioRxiv, 2025.2009.2010.675376. 10.1101/2025.09.10.675376.

66. Zarif, J.C., Hernandez, J.R., Verdone, J.E., Campbell, S.P., Drake, C.G., and Pienta, K.J. (2018). A Phased Strategy to Differentiate Human CD14+ Monocytes into Classically and Alternatively Activated Macrophages and Dendritic Cells. BioTechniques 61, 33–41. 10.2144/000114435.

67. Jorgensen, I., and Miao, E.A. (2015). Pyroptotic cell death defends against intracellular pathogens. Immunological Reviews 265, 130–142. 10.1111/imr.12287.

68. McStay, G.P., Salvesen, G.S., and Green, D.R. (2008). Overlapping cleavage motif selectivity of caspases: implications for analysis of apoptotic pathways. Cell Death & Differentiation 15, 322–331. 10.1038/sj.cdd.4402260.

69. Walsh, J.G., Cullen, S.P., Sheridan, C., Lüthi, A.U., Gerner, C., and Martin, S.J. (2008). Executioner caspase-3 and caspase-7 are functionally distinct proteases. Proceedings of the National Academy of Sciences 105, 12815–12819. doi:10.1073/pnas.0707715105.

70. Van de Craen, M., Declercq, W., Van den brande, I., Fiers, W., and Vandenabeele, P. (1999). The proteolytic procaspase activation network: an in vitro analysis. Cell death and differentiation 6, 1117–1124. 10.1038/sj.cdd.4400589.

71. Sagulenko, V., Vitak, N., Vajjhala, P.R., Vince, J.E., and Stacey, K.J. (2018). Caspase-1 Is an Apical Caspase Leading to Caspase-3 Cleavage in the AIM2 Inflammasome Response, Independent of Caspase-8. J Mol Biol 430, 238–247. 10.1016/j.jmb.2017.10.028.

72. Kang, S.J., Wang, S., Hara, H., Peterson, E.P., Namura, S., Amin-Hanjani, S., Huang, Z., Srinivasan, A., Tomaselli, K.J., Thornberry, N.A., et al. (2000). Dual role of caspase-11 in mediating activation of caspase-1 and caspase-3 under pathological conditions. J Cell Biol 149, 613–622. 10.1083/jcb.149.3.613.

73. Tsuchiya, K., Hosojima, S., Hara, H., Kushiyama, H., Mahib, M.R., Kinoshita, T., and Suda, T. (2021). Gasdermin D mediates the maturation and release of IL-1α downstream of inflammasomes. Cell Reports 34. 10.1016/j.celrep.2021.108887.

74. Planès, R., Pinilla, M., Santoni, K., Hessel, A., Passemar, C., Lay, K., Paillette, P., Valadão, A.L.C., Robinson, K.S., Bastard, P., et al. (2022). Human NLRP1 is a sensor of pathogenic coronavirus 3CL proteases in lung epithelial cells. Molecular Cell 82, 2385–2400.e2389. 10.1016/j.molcel.2022.04.033.

75. Lei, X., Zhang, Z., Xiao, X., Qi, J., He, B., and Wang, J. (2017). Enterovirus 71 Inhibits Pyroptosis through Cleavage of Gasdermin D. J Virol 91. 10.1128/jvi.01069-17.

76. Luchetti, G., Roncaioli, J.L., Chavez, R.A., Schubert, A.F., Kofoed, E.M., Reja, R., Cheung, T.K., Liang, Y., Webster, J.D., Lehoux, I., et al. (2021). Shigella ubiquitin ligase IpaH7.8 targets gasdermin D for degradation to prevent pyroptosis and enable infection. Cell Host Microbe 29, 1521–1530 e1510. 10.1016/j.chom.2021.08.010.

77. Kim, S.S., and Seo, S.R. (2024). Caspase-3 targets pro-interleukin-1beta (IL-1beta) to restrict inflammation. FEBS Lett 598, 1366–1374. 10.1002/1873-3468.14864.

78. Bibo-Verdugo, B., Snipas, S.J., Kolt, S., Poreba, M., and Salvesen, G.S. (2020). Extended subsite profiling of the pyroptosis effector protein gasdermin D reveals a region recognized by inflammatory caspase-11. The Journal of biological chemistry 295, 11292–11302. 10.1074/jbc.RA120.014259.

79. Gu, Y., Kuida, K., Tsutsui, H., Ku, G., Hsiao, K., Fleming, M.A., Hayashi, N., Higashino, K., Okamura, H., and Nakanishi, K. (1997). Activation of Interferon-γ Inducing Factor Mediated by Interleukin-1β Converting Enzyme. Science 275, 206.

80. Puri, A.W., Broz, P., Shen, A., Monack, D.M., and Bogyo, M. (2012). Caspase-1 activity is required to bypass macrophage apoptosis upon Salmonella infection. Nat Chem Biol 8, 745–747. 10.1038/nchembio.1023.

81. Kayagaki, N., Kornfeld, O.S., Lee, B.L., Stowe, I.B., O’Rourke, K., Li, Q., Sandoval, W., Yan, D., Kang, J., Xu, M., et al. (2021). NINJ1 mediates plasma membrane rupture during lytic cell death. Nature 591, 131–136. 10.1038/s41586-021-03218-7.

82. Fattinger, S.A., Maurer, L., Geiser, P., Bernard, E.M., Enz, U., Ganguillet, S., Gul, E., Kroon, S., Demarco, B., Mack, V., et al. (2023). Gasdermin D is the only Gasdermin that provides protection against acute Salmonella gut infection in mice. Proc Natl Acad Sci U S A 120, e2315503120. 10.1073/pnas.2315503120.

83. Abele, T.J., Billman, Z.P., Li, L., Harvest, C.K., Bryan, A.K., Magalski, G.R., Lopez, J.P., Larson, H.N., Yin, X.-M., and Miao, E.A. (2023). Apoptotic signaling clears engineered Salmonella in an organ-specific manner. eLife.

84. Ding, X., Kambara, H., Guo, R., Kanneganti, A., Acosta-Zaldivar, M., Li, J., Liu, F., Bei, T., Qi, W., Xie, X., et al. (2021). Inflammasome-mediated GSDMD activation facilitates escape of Candida albicans from macrophages. Nat Commun 12, 6699. 10.1038/s41467-021-27034-9.

85. Grant, S., Nicholson, E., Chen, Z.H.X., Robson, C., Viswanathan, B., Chang, Y., Sung, A.W., Patel, P., Kleppen, J., Ozdenya, T., et al. (2025). Evolutionary convergence of a Pyroptosis-Apoptosis crosstalk drives non-canonical inflammasome signalling. bioRxiv, 2025.2011.2001.685735. 10.1101/2025.11.01.685735.

86. Okondo, M.C., Johnson, D.C., Sridharan, R., Go, E.B., Chui, A.J., Wang, M.S., Poplawski, S.E., Wu, W., Liu, Y., Lai, J.H., et al. (2016). DPP8/9 inhibition induces pro-caspase-1-dependent monocyte and macrophage pyroptosis. Nature chemical biology 13, 46–53. 10.1038/nchembio.2229.

87. Helaine, S., Thompson, J.A., Watson, K.G., Liu, M., Boyle, C., and Holden, D.W. (2010). Dynamics of intracellular bacterial replication at the single cell level. Proceedings of the National Academy of Sciences 107, 3746–3751. doi:10.1073/pnas.1000041107.

