## Supplemental Figures for "Human non-canonical inflammasomes activate CASP3 to limit intracellular *Salmonella* replication in macrophages"

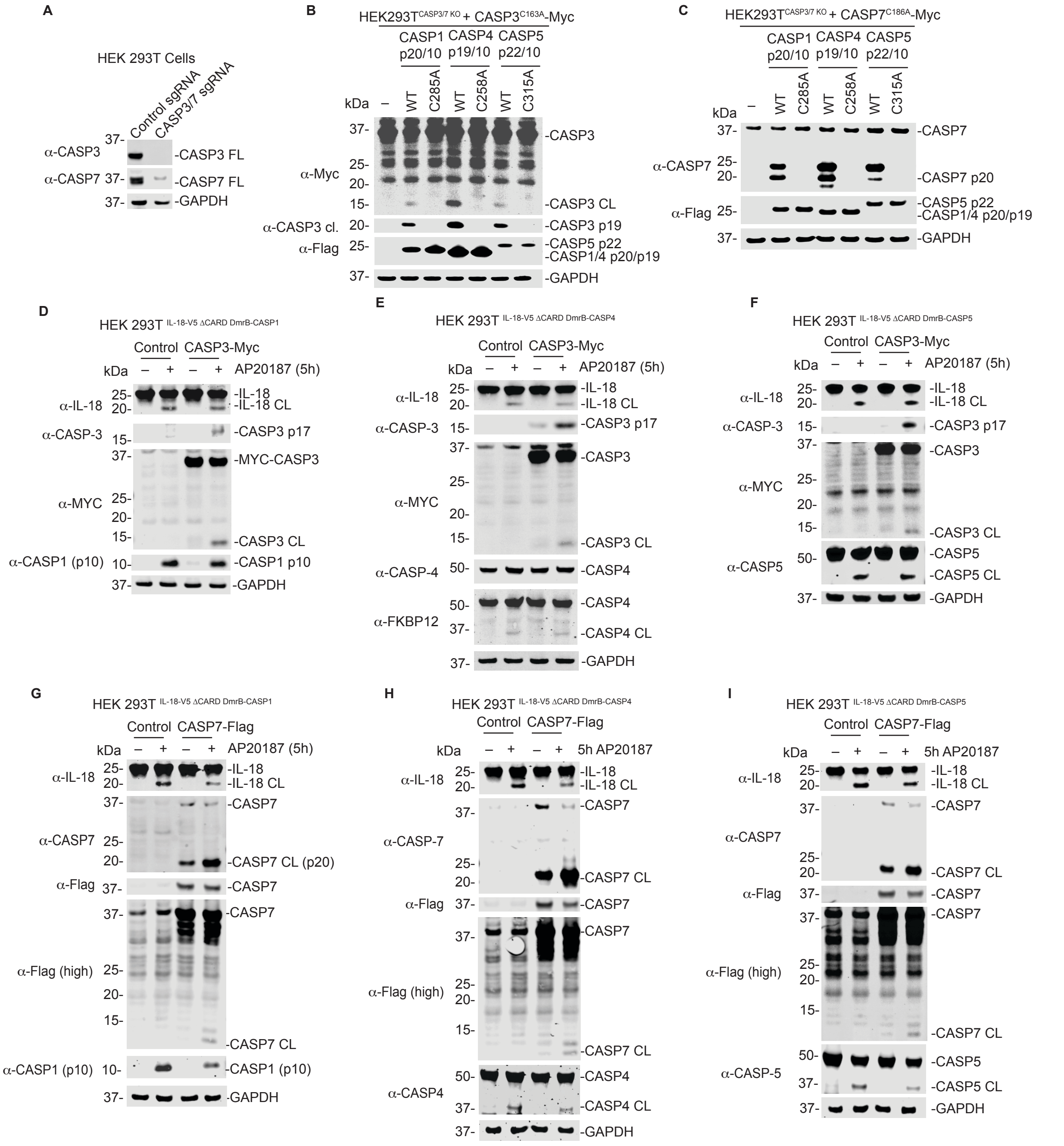


**Figure S1. CASP1, -4, and -5 cleave CASP3 and CASP7 in cells. (A)** Confirmation of CASP3/7 knockout in HEK 293T cells by immunoblotting **(B,C)** *CASP3/7* KO HEK 293T cells were transiently transfected with indicated constructs and analyzed by immunoblotting. **(D-I)** HEK 293T cells stably expressing IL-18-V5 and ΔCARD DmrB-CASP1, 4, or 5 were transiently transfected with indicated constructs coding for C-terminally Myc-tagged CASP3 (CASP3-Myc) **(D-F)** or C-terminally Flag-tagged CASP7 (CASP7-Flag) **(G-I)** for 24 h. Cells were then treated with 1 μM

AP20187 for 5 h to activate the DmrB-caspases and cell lysates were collected and assessed by immunoblotting. Data are representative of three or more independent experiments.

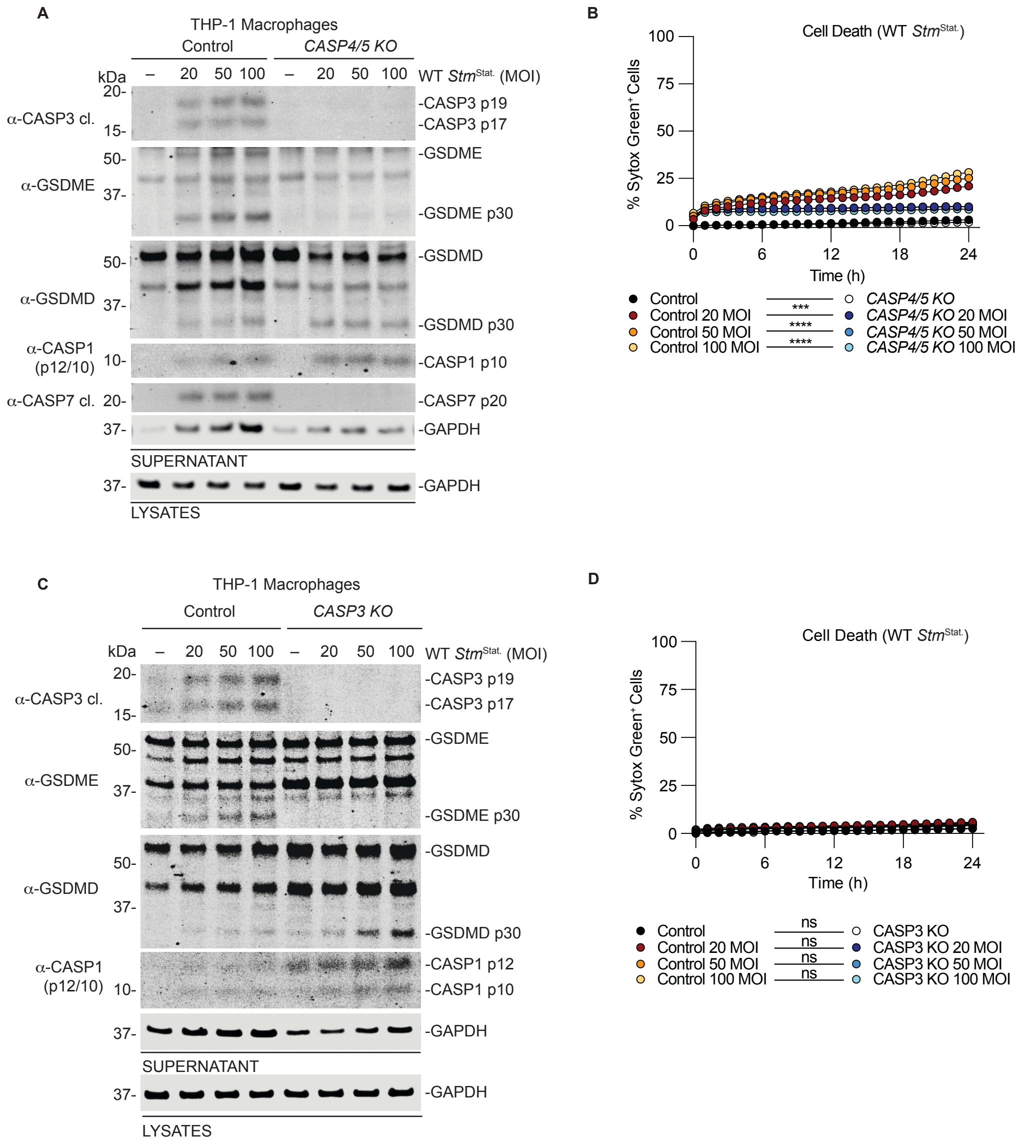


**Figure S2. Stationary phase Salmonella induces modest CASP4/5 and CASP3-dependent activation of pyroptosis.** (**A,B**) WT and *CASP4/5* KO THP-1 macrophages were treated with the indicated MOI of stationary phase *Salmonella* for 24 h then supernatants and lysates were analyzed by immunoblotting (**A**), and cell death was measured by monitoring Sytox Green uptake (**B**). (**C,D**) WT and *CASP3* KO THP-1 macrophages were treated with the indicated MOI of stationary phase *Salmonella* for 24 h then supernatants and lysates were analyzed by immunoblotting (**C**), and cell death was measured by monitoring Sytox Green uptake (**D**). Data are means ± SEM of three independent experiments. ****$P < 0.0001$, ***$P < 0.001$, **$P < 0.01$, and *$P < 0.05$ by two-way ANOVA test with Tukey's multiple comparison test comparing the control treated to KO treated samples at 24 h.
